# Evaluation of cell segmentation methods without reference segmentations

**DOI:** 10.1101/2021.09.17.460800

**Authors:** Haoran Chen, Robert F. Murphy

## Abstract

Cell segmentation is a cornerstone of many bioimage informatics studies and inaccurate segmentation introduces error in downstream analysis. Evaluating segmentation results is thus a necessary step for developing segmentation methods as well as for choosing the most appropriate method for a particular type of sample. The evaluation process has typically involved comparison of segmentations to those generated by humans, which can be expensive and subject to unknown bias. We present here an approach to evaluating cell segmentation methods without relying upon comparison to results from humans. For this, we defined a number of segmentation quality metrics that can be applied to multichannel fluorescence images. We calculated these metrics for 14 previously-described segmentation methods applied to datasets from 4 multiplexed microscope modalities covering 5 tissues. Using principal component analysis to combine the metrics we defined an overall cell segmentation quality score and ranked the segmentation methods. We found that two deep learning-based methods performed the best overall, but that results for all methods could be significantly improved by postprocessing to ensure proper matching of cell and nuclear masks. Our evaluation tool is available as open source and all code and data are available in a Reproducible Research Archive.

## Introduction

Cell segmentation is the task of defining cell boundaries in images. It is a fundamental step for many image-based cellular studies, including analysis and modeling of subcellular patterns (Boland & Murphy, 2001), analysis of changes upon various perturbations (Carpenter et al., 2006), cell tracking for investigating cell migration and proliferation (Garay et al., 2013), and cell morphology analysis for discovering cell physiological states (Rittscher, 2010). Inaccurate cell segmentation introduces potential systematic error in all these downstream analyses. Since different methods may perform differently for different imaging modalities or tissues, it is important to evaluate potential cell segmentation methods to choose the most suitable one for a specific application.

Existing segmentation methods can be divided into two categories, the geometry-based segmentation techniques (i.e. traditional computer vision techniques) and deep learning-based approaches. The former includes but is not limited to threshold-based segmentation (Shen et al., 2018), region-based segmentation (Panagiotakis & Argyros, 2018), watershed algorithm and its variants (Ji et al., 2015), active contour (Wu et al., 2015), Chan-Vese segmentation (Braiki et al., 2020; Fan et al., 2013), and Graph-cut based segmentation (Oyebode & Tapamo, 2016). In the deep learning category, the conventional deep convolutional neural network (CNN) was initially applied on various cell segmentation tasks (Jung et al., 2019; Sadanandan et al., 2017). CNN models learn the feature mapping of an image and convert the feature map into a vector for pixelwise classification (i.e. segmentation). Since its publication, the U-Net model and its variants have become a widely used alternative (Al-Kofahi et al., 2018; Falk et al., 2019; Long, 2020; Ronneberger et al., 2015). Instance-segmentation methods such as Mask R-CNN, in addition, are prominently utilized for cell and nucleus segmentation. (Fujita & Han, 2020; Johnson, 2018; Lv et al., 2019) Ensemble methods with multiple deep learning frameworks were also developed specifically for nuclei segmentation (Kablan et al., 2020; Vuola et al., 2019).

The typical approach for evaluating cell segmentation methods is to compare the segmentation results with human-created segmentation masks. For example, early work (Bamford, 2003) evaluated geometry-based algorithms on 20,000 cellular images annotated by three independent nonexpert observers. The author argued that evaluating cell segmentation does not necessarily require expert annotation. Caicedo et al. (Caicedo et al., 2019) evaluated multiple deep learning strategies for cell/nuclei segmentation by calculating the accuracy and types of errors in contrast to the expert annotations. The authors created a prototype annotation tool to facilitate the annotation process. At the model development level, deep learning-based segmentation methods design loss functions to evaluate the similarity between the current segmentation mask and the human-created mask (Al-Kofahi et al., 2018; Kromp et al., 2020), which makes the model mimic the logic of human segmentation. Unfortunately, comparison to human-created segmentations assumes that people are good at this task; however, human segmentation can suffer from extensive intra and inter-observer variability (Vicar et al., 2019; Wiesmann et al., 2017). In addition, human segmentation can require extensive labor and cost. For example, the development of the latest DeepCell version (Greenwald et al., 2022) required a very large number of human evaluations (and would have required far more if an active machine learning approach had not been used).

An alternative approach is creating simulated cell images (Wiesmann et al., 2017) to assess segmentation performances. The simulated images along with their segmentation results are generated based on various information obtained from real fluorescent images including cell shape, cell texture, and cell arrangement. Since the image is simulated, the correct segmentation is known. Even though this method is efficient and reproducible, it reaches its limits while encountering images with high cell shape and texture variability.

For this study, we sought to define an objective evaluation approach that does not require a reference segmentation. We were motivated in large part by the desire to optimize the pipeline used for analysis of multichannel tissue images as part of the Human BioMolecular Atlas Program (HuBMAP) (HuBMAPConsortium, 2019). We first defined a series of metrics based upon assumptions about the desired characteristics of good cell segmentation methods. We then identified currently available cell segmentation methods that had pre-trained models and evaluated their performance on many images from multichannel imaging modalities. A Principal Component Analysis (PCA) model was then trained using the metrics computed from the segmentation results of 11 methods on 637 multichannel tissue images across 4 imaging modalities and used to generate overall segmentation quality scores (Figure 1). We also evaluated the robustness of each method to various image degradations, such as adding noise. We found that as distributed, the Cellpose model (Stringer et al., 2021) gave the best results, but that after postprocessing to ensure proper matching of cell and nuclear masks, two different DeepCell models (Bannon et al., 2021) performed the best. We also found that our evaluation metrics are sensitive to under-segmentation error which is very common in practice. Lastly, we found that our quality scores, which are obtained without the help of any human reference, not only capture the inter-observer variance between two human experts, but also have high correlation with three cell segmentation benchmarks using expert annotations.

**Figure 1.**
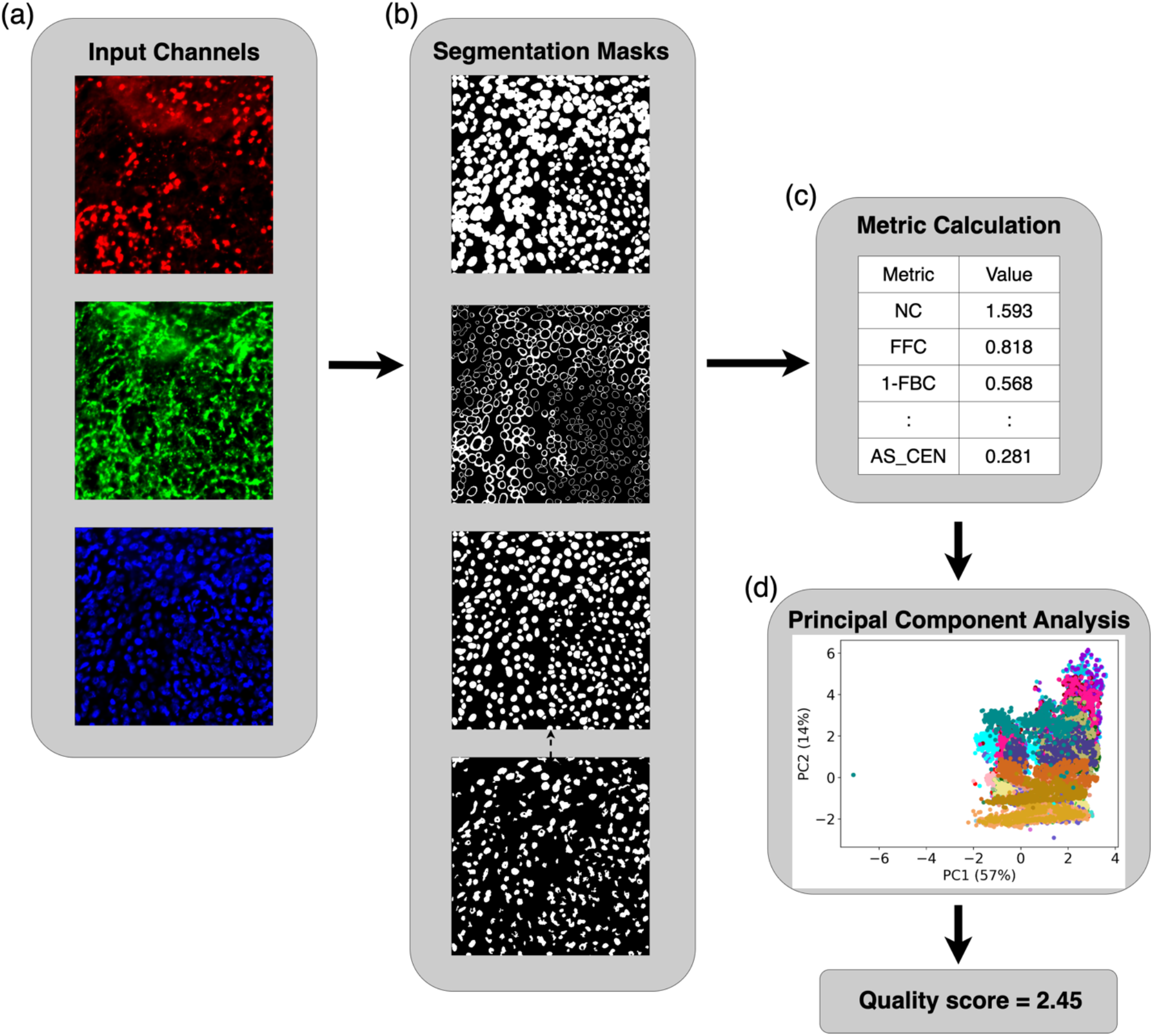
Our pipeline for cell segmentation evaluation. (a) Input channels for cell membrane (red), cytoplasm (green), and nucleus (blue) are available to each segmentation method, but methods only used one or two channels. (b) Methods generate segmentation masks: cell mask, cell outside nucleus mask, and nuclear mask (shown in the top three panels, respectively). For segmentation methods that don’t output the nuclear mask, we used an Otsu-thresholded nuclear mask (bottom panel) as a substitute. We removed the pixels in the nuclear mask that were outside the cell membrane in the cell mask, as well as the cells and nuclei that didn’t have the corresponding nuclei and cells. (c) For each set of segmentation masks, we calculated 14 metrics to objectively evaluate the quality of segmentation under various assumptions. We then applied Principal Component Analysis on the matrix of all metrics for all methods and all images. (d) The scatter plot shows a point for each segmentation for each image. Different colors represent different segmentation methods. Finally, a variance-weighted sum of PC1 and PC2 was used to generate an overall quality score for each combination of method and image.

To enable use by other investigators, we provide open source software that is able to evaluate the performance of any cell segmentation method on multichannel image inputs.

## Methods

### Images

We obtained image data from the HuBMAP project, for 4 multiplexed modalities. For each modality, the nuclear, cytoplasmic, and cell membrane channels chosen for segmentation (Supplementary Table 1) were either those recommended by the Tissue Mapping Center (TMC) that produced the datasets, or selected based on peer-reviewed literature and subcellular localization of antibody targeted protein annotated by UniProt or Gene Ontology. Details about input markers are summarized in Supplementary Table 2.

#### CODEX

CO-Detection by indEXing (CODEX) is an imaging technique generating highly multiplexed images of fluorescently tagged antigens (Goltsev et al., 2018). The HuBMAP portal (https://portal.hubmapconsortium.org/) contains 26 CODEX datasets over 5 tissues (large intestine, small intestine, thymus, spleen, and lymph node) produced by two TMCs: Stanford University and the University of Florida. The datasets from the Stanford TMC all contain 47 channels. Each dataset consists of 4 tissue regions, which are individually split into a grid of tiles with ~1440 x 1920 size. The University of Florida TMC generated 11-channel CODEX datasets. Each dataset only has one region which is divided into a grid in the same manner. To ensure both tissue coverage and evaluation efficiency, we created evaluation datasets with a subset of the tiles chosen to be non-neighboring (to avoid evaluating cells in the overlap region twice) and tiles that were not on the edge of the grid (to prevent large fractions of the image consisting of background on the edge of tissue). 393 tiles in total were selected. The pixel size of CODEX images is 0.37745 micron.

The different antigens used by the two TMCs required us to select different channels as segmentation input. For Stanford TMC datasets, we used Hoechst, Cytokeratin, and CD45 channels as nuclear, cytoplasmic, and cell membrane channels, respectively. For the University of Florida TMC datasets, we used DAPI, CD107a, and E-cadherin. To ensure consistency of evaluation results from CODEX data, only the 5 channels that are common between the datasets of the two TMCs were used to calculate channel homogeneity metrics (see Supplementary Methods).

#### Cell DIVE

Cell DIVE is another antibody-based multiplexing technique (Gerdes et al., 2013). We had access to 12 regions of Cell DIVE data. Each region contains 26 images with 19 channels. Cell DIVE datasets are much larger than those from CODEX, consisting of ~10000 x ~15000 pixels. To boost the efficiency of the pipeline while ensuring the coverage of datasets, we took the first image of each region and split each image into a grid of tiles similar to those of CODEX datasets. Each tile has roughly 1000 x 1000 pixels which is efficiently applicable to all segmentation methods. To exclude tiles with few cells, we calculated three metrics on each channel of each tile and applied KMeans clustering to identify a cluster consisting of images with dense cells (see Supplementary Figure 1). We selected DAPI, Cytokeratin, and P-cadherin as nuclear, cytoplasmic, and cell membrane inputs for segmentation. The pixel size of Cell DIVE images is 0.325 micron.

#### MIBI

MIBI (Multiplexed Ion Beam Imaging) uses a Secondary-Ion Mass Spectrometer (SIMS) to image antibodies tagged with monoisotopic metallic reporters (Angelo et al., 2014). We obtained 10 29-channel MIBI datasets. Each image consists of 1024 x 1024 pixels. HH3, PanKeratin, and E-cadherin were utilized as the nuclear, cytoplasmic, and cell membrane channels for segmentation. The spot size (equivalent to pixel size) of MIBI images is 0.391 micron.

#### IMC

As an expansion of Mass Cytometry, Imaging Mass Cytometry (IMC) uses laser ablation to generate plumes of particles that are carried to the mass cytometer by a stream of inert gas (Chang et al., 2017). 13 in total of 2D 39-channel IMC datasets of spleen, thymus, and lymph node were available from the University of Florida TMC. In this modality, lr191, SMA, and HLA-ABC were chosen as input channels for segmentation. For images without lr191 channel, Histone channel was substituted as nuclear input. The pixel size of IMC images is 1 micron.

### Segmentation Methods

We created a python wrapper for each method to adapt all methods to a common pipeline. The input channels required by each method are shown in Supplementary Table 1. All methods generate a segmentation mask for cell boundaries in an indexed image format with some methods also generating nuclear boundaries.

#### DeepCell

DeepCell is designed for robustly segmenting mammalian cells (Bannon et al., 2021; Greenwald et al., 2022; Moen et al., 2019; Van Valen et al., 2016). A feature pyramid network (Lin et al., 2017) with PanopticNets architecture trained by greater than 1 million paired whole-cell and nuclear annotations is embedded in the latest deepcell-tf package on the Python TensorFlow platform. We tested three versions of DeepCell software with different trained deep learning models. For each model, a nuclear intensity channel is a mandatory input and we chose cytoplasm or cell membrane as the secondary input image to generate two different whole-cell segmentation masks (these were treated as different methods). The latest version of DeepCell (v0.12.3) requires specification of pixel size in microns.

#### Cellpose

Cellpose is a generalist, U-Net-based algorithm that segments cells in various image modalities (Stringer et al., 2021). The key idea of Cellpose is to track the gradient flow in the labels to predict cell boundaries. A PyTorch model “cyto” was trained with multiple datasets mainly with cytoplasmic and nuclear markers. We applied three versions of Cellpose with different pre-trained models to segment whole cells on 2D cytoplasmic and nuclear intensity images. Unlike DeepCell, which uses the same model to generate whole-cell and nuclear masks, Cellpose contains a separate U-Net “nuclei” model for the segmentation of nuclear channels.

#### CellProfiler

CellProfiler 4.0 (Carpenter et al., 2006; Kamentsky et al., 2011) contains a module for cell segmentation that applies traditional thresholding and propagation algorithms to segment nuclei and then cells. Required parameters that define the range of acceptable nuclear sizes and cytoplasmic thickness were chosen for each imaging dataset. To facilitate batch computation and avoid CellProfiler’s 8-bit cell index limitation in saving output images, we modified the CellProfiler module to directly store the indexed segmentation masks as NumPy arrays.

#### CellX

CellX is a MATLAB package that using traditional image processing operations and requires only a membrane marker image as input (Dimopoulos et al., 2014; Mayer et al., 2013). Required parameters that define the range of acceptable cell radii and maximum cell length (since cells are not perfect circle) were chosen for each dataset. Since the algorithm does not generate a nuclear mask, we paired the cell mask with a default Otsu-thresholded nuclear mask (see below under Voronoi segmentation).

#### Cellsegm

Cellsegm is a MATLAB-based toolbox providing automated whole-cell segmentation (Hodneland et al., 2013). Required parameters that define the range of acceptable cell sizes were chosen for each dataset. Since the algorithm does not generate a nuclear mask, we paired the cell mask with a default Otsu-thresholded nuclear mask.

#### The Allen Cell Structure Segmenter

There are two branches of the Allen Cell Structure Segmenter (AICS): a deep learning version and a classic image processing version (Chen et al., 2020). The deep learning version of AICS only provides a nuclear segmentation so it was not evaluated. The classic version consists of thresholding and filtering operations and requires estimated minimum cell areas (which was chosen for each dataset based on pixel size). Since the algorithm does not generate a nuclear mask, we paired the cell mask with a default Otsu-thresholded nuclear mask.

#### Voronoi segmentation

To provide a baseline method, we used a simple method based on the Voronoi diagram. An Otsu thresholding was applied on the nucleus channel as a first step to obtain a nuclear mask. Then a Voronoi diagram was created to partition the image along lines equidistant from the centroid of each nucleus.

### Degrading images for robustness analysis

To evaluate the robustness of each method, each image was degraded in two ways. In the first, zero-mean Gaussian noise was added to each pixel. The standard deviation was set to various levels based on the typical channel intensity of a given modality: 500, 1000, and 1500 for CODEX and Cell DIVE images, and 5, 10, and 15 for MIBI and IMC data. For the second perturbation, we downsampled the images to 70%, 50%, and 30%, respectively, on both dimensions. Note that the Gaussian noise was only added to the images used for segmentation; the evaluation of the resulting masks was done using the original images.

### Mask processing

For some segmentation methods, finding cell boundaries is done independently of finding nuclear boundaries. This may mean that the final segmentation masks include nuclei that do not have a corresponding cell boundary and vice versa. We assumed that a good segmentation method would minimize this and therefore defined a metric to capture this aspect of a segmentation method. To calculate it, each cell was matched to any nuclei contained within it. All cells that did not have corresponding nuclei were counted as mismatched, and vice versa. For cells that had multiple corresponding nuclei, the one with the smallest fraction of mismatched pixels was kept. All mismatched cells and nuclei were removed from the calculation of other metrics (see Supplementary Figure 2).

Segmentation methods may also generate misshaped nuclei that have pixels outside their corresponding cells; that is, nuclei that protrude through the cell membrane. Across all methods in all images, this occurred for an average of 60.6% of segmented nuclei. To solve this issue, we applied two posterior approaches. The first approach considered all misshaped nuclei and their corresponding cells to be mismatched even if they have a one-to-one relationship. Alternatively, we developed a “repair” pipeline that trimmed the mask of misshaped nuclei. The combination of a segmentation method followed by repair was evaluated as a distinct segmentation method.

To evaluate the segmentation performance using the different channel intensities, we expected that nuclear protein composition would be considerably different than the cytoplasm and the cell membrane. We therefore calculated our metrics using two masks: the (repaired) nuclear mask and a “Cell Excluding Nucleus” mask calculated by removing the nuclear mask from the cell mask.

After this mask processing step, each of cells has one-on-one matching relationship with its cell, nucleus and cell excluding nucleus mask.

### Evaluation metrics not requiring reference segmentation

We defined 14 metrics to evaluate the performance of a single segmentation method without requiring a reference segmentation. These are of two types: metrics that assess the coverage of a segmentation mask on the image, and metrics that measure various types of uniformity at the pixel and cell levels on multiplexed images. Each metric is derived under an assumption based on general concepts from cell biology. They are described in the Supplementary Methods and summarized in Table 2.

We also defined 10 metrics for comparing two segmentation methods (or one method with a human-generated segmentation). These are also described in the Supplementary Methods.

### Benchmarks for comparing segmentation with annotations

We adapted three benchmarks to quantify the segmentation quality comparing with expert annotations. F1 score has been widely used for benchmarking cell segmentation performance (Caicedo et al., 2019; Greenwald et al., 2022). The first step to calculate F1 score is to determine matched pairs of cells from two masks. For each cell on the reference mask (could be either mask), we calculated the Jaccard index (as equation below) with overlapping cells in the other mask.

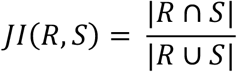

We set *JI* = 0.3 as the threshold that two cells must satisfy to be considered matched. If multiple cells have *JI* above the threshold, the one with the highest *JI* is selected. Based on this, we counted the number of True Positive cells (TP, if two cells have *JI* above threshold) as well as the False Positive and False Negative cells (FP and FN, if two cells have *JI* below threshold). And we calculated F1 score by following equations. Note that F1 score remains the same regardless which mask is the reference (swapping FP and FN won’t change F1 score). Therefore, it’s a symmetric measurement.

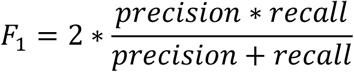

where

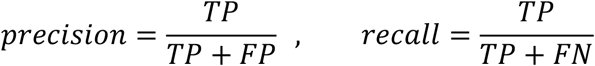

One issue of F1 score is that it demands manual selection of the *JI* threshold. Different threshold choices may lead to different conclusions. To improve this, we created a Precision-Recall curve with 0.01 threshold intervals, and integrated the area under the curve (PRAUC) as our second benchmark.

The third benchmark, SEG score, is widely applied for cell tracking (Maska et al., 2014). The original SEG score scans each cell in the reference mask and finds the best matched cell in the query mask by the following condition:

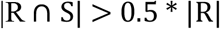

Where R is a reference cell and S is the segmented cell from a segmentation method. If the intersection (overlap) is greater than half of area of the reference cell, the two cells are considered matched. (Note that the constant 0.5 is analogous but not identical to setting a JI threshold.) If there are multiple segmented cells satisfiy this condition, the one with the highest overlap with R remains. For each refenence cell, *JI* is used to qualitify the similarity with its matched cell. For the ones with no matched cell, *JI* = 0. The final SEG score is the average of *JI* over all reference cells. One major issue with the original SEG score definition comes from its asymmetry. If the query mask contains many extra cells that do no match to any cell in the reference mask, the SEG score will be the same as if the query mask did not have any extra cells. However, segmenting extra unmatched cells is clearly worse. To solve this issue, we calculated two SEG scores, one using the human labeled mask as reference and another using the computer-generated mask as reference, and averaged them as the final benchmark. We refer to this as the SEG’ (SEG prime) score which is symmetric and accounts properly for unequal numbers of cells between the masks (and does not require an assumption of which mask is correct).

### Availability

All code needed to calculate the evaluation metrics, as well as scripts to download test images, are available at https://github.com/murphygroup/CellSegmentationEvaluator. A Reproducible Research Archive containing all code and intermediate results (which enable recreation of all figures and tables) is available at https://github.com/murphygroup/ChenMurphy2DSegEvalRRA.

## Results

### Generating masks and calculating metrics

We began by running all methods on images from four imaging modalities: CODEX, Cell DIVE, MIBI, and IMC (Supplementary Table 1). There were 637 multi-channel images in total. Each method generated whole-cell masks, and some methods also generate nuclear masks. For those methods that did not generate a nuclear mask, we provided a simple mask based on Otsu thresholding of the nuclear channel. For each method, an additional pair of masks was created with our “repair” procedure to eliminate unmatched nuclear and cell masks and remove nuclear regions outside the corresponding cell mask (see **Mask processing** in the Methods). This was done because unmatched cells and nuclei would be penalized by our evaluation metrics; the combination of the original method and the repair procedure was treated as a separate method to allow evaluation of each method either as originally provided or as most suitable for cell quantitation.

We then ran the evaluation pipeline to calculate the evaluation metrics (see Table 2 and Supplementary Methods) for each method for each image. This process yielded two matrices of 637 images x 14 methods x 14 metrics, one for the original methods and one for the methods with repair.

To examine how the individual metrics varied across methods, we performed z-score standardization on each metric and averaged the metrics over all images for each method (Figure 2). We observed that the 14 methods have heterogeneous performance on the different metrics with either non-repair and repair approaches. Methods after repair tend to have a significant increase in the average metric values (Figure 2a vs Figure 2b), and the improvement of the FMCN metric directly reflects significantly better cell-nuclei matching after repair. We noticed that methods before repair have slightly higher cell uniformity (reflected by higher 1/(ACVC+1), FPCC and AS metrics) due to a much smaller cell number. In both non-repair and repair figures, the patterns of curves for all methods except the Voronoi are similar, with curves of deep learningbased methods similar to each other. We also noticed different methods sometimes tradeoff a decrease in one or more metrics for an increase in others. For instance, DeepCell 0.6.0 and DeepCell 0.9.0 have opposite behavior on the NC and FFC metrics in Figure 2b. This comes from the fact that despite similar deep learning algorithms applied, DeepCell 0.6.0 tends to segment fewer but larger cells than DeepCell 0.9.0. Our metrics accurately reflect this phenomenon (relatively lower NC but higher FFC). Our metrics also captured the improvement of DeepCell 0.12.3, which has a much larger training dataset than two previous versions, with both relatively high NC and FFC. We also observed that different metrics show different range of variance for the 14 methods. For instance, there is large variation among methods with respect to NC metric (number of cells per 100 squared microns) and FFC (fraction of image foreground occupied by cells) but less variation in 1/(ACVC_NUC+1) values (average of weighted average CV of cell type intensities over 1-10 clusters on nuclei).

**Figure 2.**
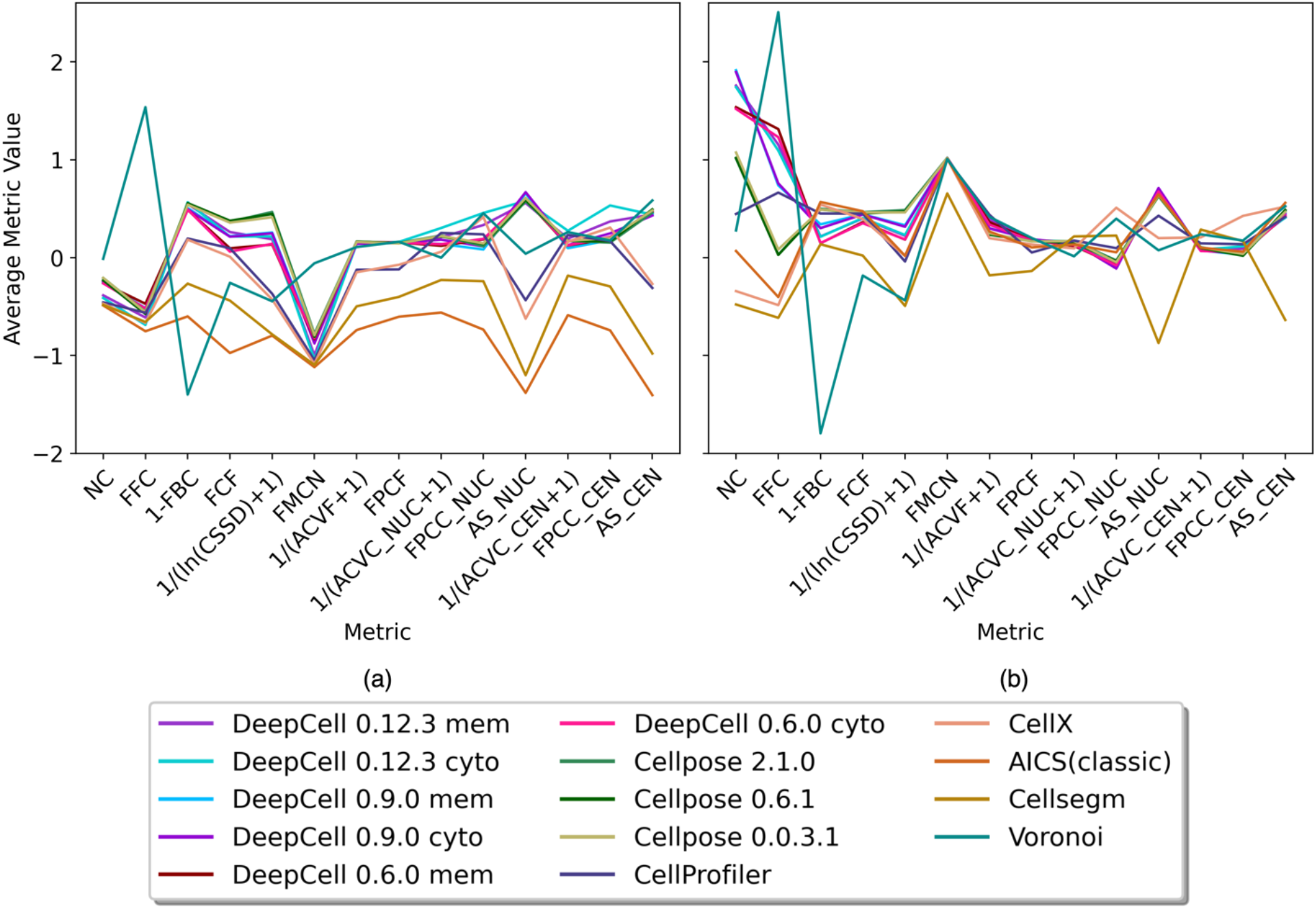
Heterogeneity of methods performance. Each metric was z-scored. For each segmentation method, each metric was averaged across all images. Panel (a) shows results from methods evaluated without mask repair (see text). Panel (b) shows the results from methods after the repair.

### Evaluating sensitivity of methods to added noise or downsampling

To test the robustness of the methods, we created perturbed images with various amounts of added zero-mean Gaussian noise or various extents of downsampling (see Methods) and evaluated the quality of the resulting segmentations. The Gaussian perturbations were done only to the images provided to the segmentation method, and not to the multichannel images used to calculate the metrics. The downsampling was also done on the multichannel images, matching the same size as segmentation masks to calculate the metrics. This allowed us to assess method robustness: how sensitive the results for a given segmentation method were to the quality of the image. It also provided a parallel check of how well our metrics were performing based on the assumption that a perturbed image should yield a worse segmentation.

### Segmentation quality score

One of our primary goals was to provide an overall segmentation quality score for each method on each image. To do this we created a Principal Component Analysis (PCA) model using the metrics for all methods for all images with and without perturbation. The metrics were z-scored before performing PCA. The top 2 principal components (PCs) for each method across all modalities and images are shown in Figure 3 (and Supplementary Figure 3) with and without various amounts of perturbation. Since the number of datasets available for each modality varied, we averaged the top 2 PCs across all images within each modality and then averaged across the modalities to balance the contributions from all modalities to the final model. With random Gaussian perturbation (Figure 3a and Supplementary Figure 3a), we observed that PC1 values for all methods tend to decrease as perturbation increases, confirming that the metrics perform as anticipated. We also observed that most of methods show decreased PC1 with increased downsampling (Figure 3b and Supplementary Figure 3b), with some exceptions. The exceptions occur because the downsampling process also removed noise in the images, causing some methods to get a slightly better PC1 score after downsampling. Thus, PC1 alone is not sufficient as an overall quality score.

**Figure 3.**
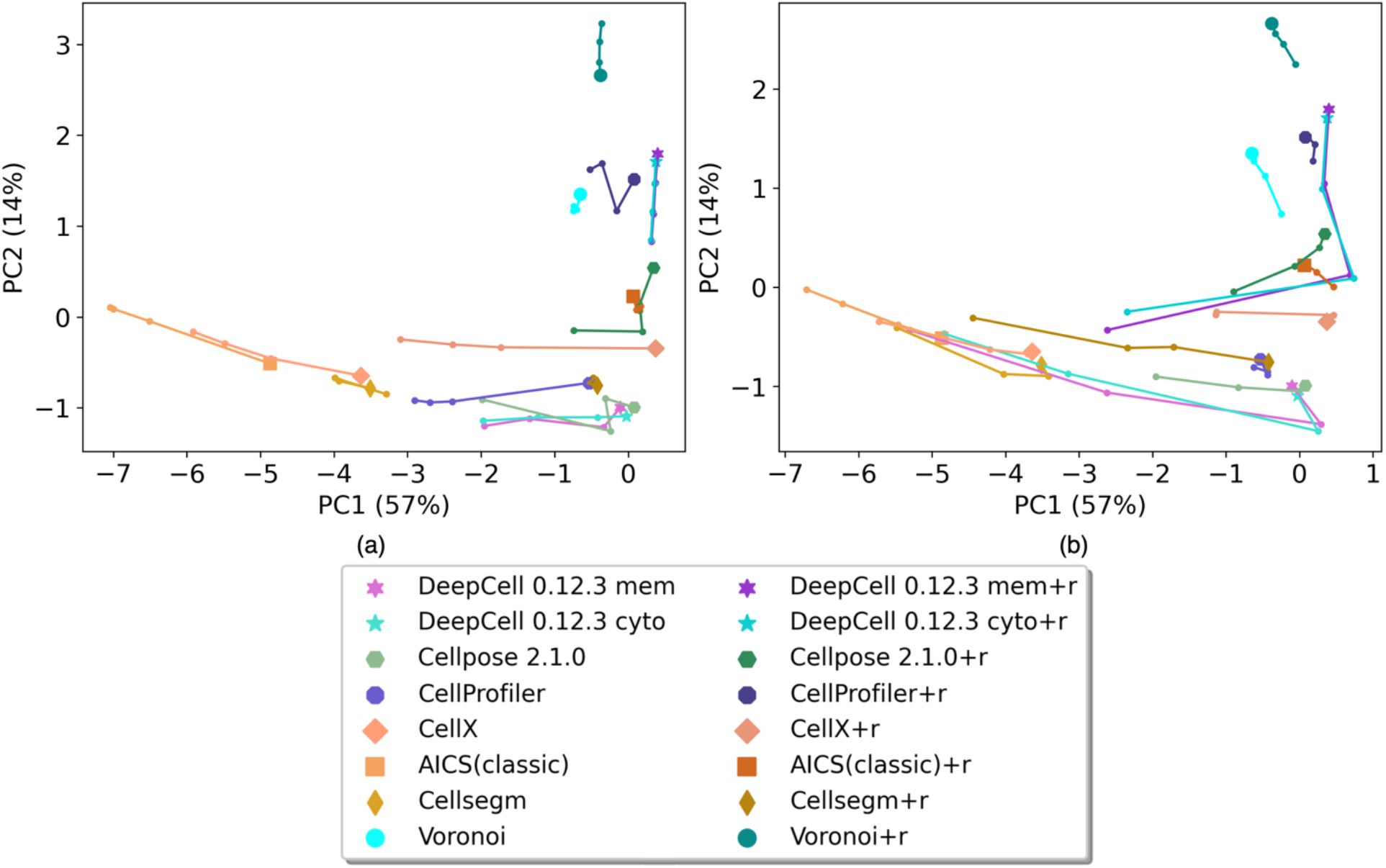
Top 2 Principal components of each method on images from all modalities. (a) Results from images with Gaussian noise perturbation. (b) Result from images with downsampling. Each method is represented by a unique color. The points with a unique marker shape represent unperturbed images. The trajectories represent on low, medium and high Gaussian noise perturbation (a), or small, medium, large degrees of downsampling (b). Only the results from the latest DeepCell and Cellpose are shown for better visualization. Supplementary Figure 3 shows results from the earlier versions.

The loadings of PC1 (Supplementary Figure 4a) are all positive showing that all metrics have a synergistic effect on PC1 values. On the other hand, PC2 loadings (Supplementary Figure 4b) show that it is primarily an indicator of the overall coverage of a mask with high NC and FFC loadings. We therefore adopted the sum of PC1 and PC2 weighted by their explained variance as our final overall quality score. Rankings based on the quality score for all methods (with and without repair) averaged across all modalities and (unperturbed) images are shown in Figure 4, with Voronoi segmentation as a reference baseline. As expected, repair generally improves the overall metric for most methods. We observed that DeepCell methods have the highest overall performance with repair, and that they are also robust to both random Gaussian noise and downsampling (Figure 3). Cellpose methods perform the best among methods without repair. CellProfiler scores the best among methods with non-deep learning algorithms. Methods (e.g. CellX, AICS classic) that are primarily used for cultured cell segmentation (rather than tissue segmentation) tend to perform worse than the Voronoi baseline.

**Figure 4.**
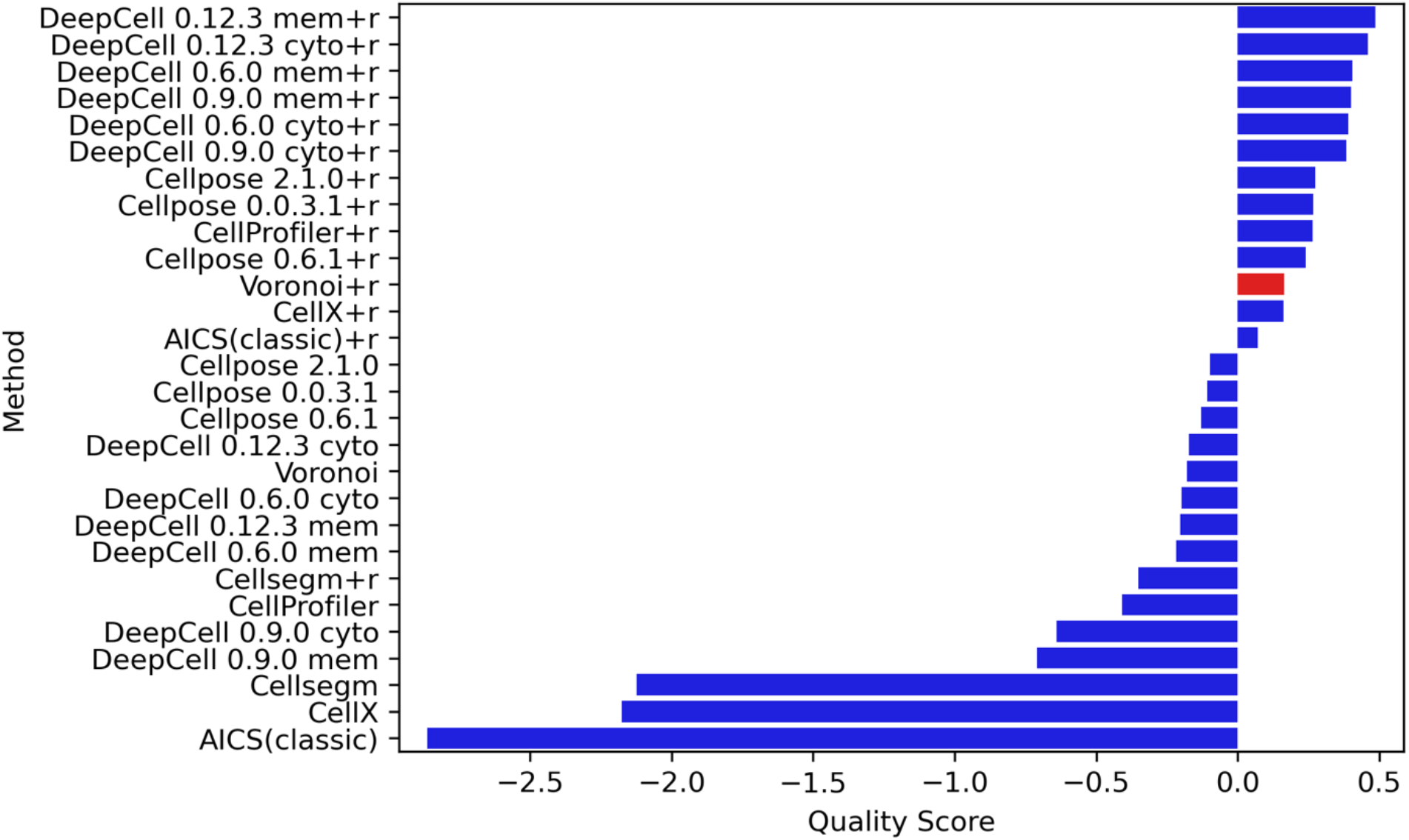
Overall quality score rankings of all modalities. All methods with and without repair are ranked by their quality scores. The methods that have a higher score than Voronoi are considered acceptable.

### Model selection for particular tissue and data modality

One of the major potential uses of our evaluation pipeline is the potential ability to select the most appropriate method for a specific imaging modality or tissue. Images from different modalities have a large variation in signal-to-noise and spatial resolution that may influence the performance of segmentation methods. We therefore separately evaluated images from different modalities (Supplementary Figures 5-8). While it performed the best across different modalities, the version of DeepCell that was optimal varied. Perhaps because they had the lowest resolution among the four modalities (1 micron pixel size), the relative ranking of the methods was quite different for IMC images compared to other modalities. We also observed that CellProfiler ranked in the middle of DeepCell methods for CellDIVE and IMC modalities.

Segmentation method performance may also be expected to vary for different tissues due to potential differences in the shapes and spatial arrangement of cells. We therefore separately analyzed performance on five tissues: small intestine, large intestine, spleen, thymus, and lymph node (images for the first two tissues are solely from CODEX and the latter three are available for both CODEX and IMC) (Supplementary Figures 9-13). We observed that while DeepCell models still performed the best among all methods (with the latest version being optimal), the rankings and overall quality scores of methods on different tissues vary. Interestingly, strong, similar performance was observed for both DeepCell and Cellpose on large intestine.

### Evaluating expert annotated segmentation

Since previous cell segmentation studies have focused on comparison with human experts, we used two lymph node CODEX multi-channel images (tiles R001_X003_Y004 and R001_X004_Y003 in dataset HBM279.TQRS.775) and accompanying cell segmentation masks annotated by two experts (Dayao et al., 2022) to compare performance of various segmentation methods. We processed those images using all segmentation methods listed in Table 1 and evaluated the segmentation outputs along with expert-annotated masks. Since the expert annotations did not include nuclear masks, we slightly modified our evaluation metrics to calculate the cell uniformity metrics (i.e. 1/(ACVC +1), FPCC and AS in Table 2) only on the cell masks and removed the FMCN metric. Accordingly, we retrained the PCA model (using all datasets) with the reduced number of metrics (10 instead of 14) for use only in this comparison with expert annotations.

**Table 1.** Segmentation methods evaluated

| Method | Inputs |  |  |  | Output |
| --- | --- | --- | --- | --- | --- |
|  | Cytoplasm | Cell Membrane | Nucleus | Requires parameters<sup>1</sup> | Nuclear mask |
| DeepCell 0.12.3 |  | X | X | Yes | Yes |
| DeepCell 0.12.3 | X |  | X | Yes | Yes |
| DeepCell 0.9.0 |  | X | X | No | Yes |
| DeepCell 0.9.0 | X |  | X | No | Yes |
| DeepCell 0.6.0 |  | X | X | No | Yes |
| DeepCell 0.6.0 | X |  | X | No | Yes |
| Cellpose 2.1.0 | X |  | X | No | Yes |
| Cellpose 0.6.1 | X |  | X | No | Yes |
| Cellpose 0.0.3.1 | X |  | X | No | Yes |
| CellSegm |  | X | X | Yes | No |
| CellX |  | X |  | Yes | No |
| CellProfiler |  | X | X | Yes | Yes |
| AICS (classic) |  | X | X | Yes | No |
| Voronoi |  |  | X | No | Yes |

<sup>1</sup> The required parameters define the range of acceptable cell sizes.

**Table 2.** Summary of segmentation metrics

| Name | Metric | Mask(s) to calculate the metrics on |
| --- | --- | --- |
| Number of cells per 100 squared microns | NC | Matched cell mask |
| Fraction of image foreground occupied by cells | FFC | Matched cell mask |
| Fraction of image background occupied by cells | 1-FBC | Matched cell mask |
| Fraction of cell mask in foreground | FCF | Matched cell mask |
| Fraction of match between cells and nuclei | FMCN | Cell and nuclear masks prior to mask processing |
| Average CV of foreground pixels outside the cells | $1/(ACVF+1)$ | Matched cell mask |
| Fraction of first PC of foreground pixels outside the cells | FPCF | Matched cell mask |
| Average of weighted average CV of cell type intensities over 1-10 clusters | $1/(ACVC+1)$ | Matched nuclear mask (NUC) and cell excluding nucleus mask (CEN) |
| Average of weighted average fraction of the first PC of cell type intensities over 1-10 clusters | FPCC | Matched nuclear mask (NUC) and cell excluding nucleus mask (CEN) |
| Average of Silhouette score of clustering over 2-10 clusters | AS | Matched nuclear mask (NUC) and cell excluding nucleus mask (CEN) |
| Standard deviation of cell size | $1/(\ln(CSSD)+1)$ | Matched cell mask |

The quality score ranking and top 2 PCs plot in Figure 5a reflects relatively high accuracy expert annotation and also reveals the inter-observer variability we describe in the Introduction. While Expert1 is among the best in the ranking, Expert2 has an overall score below the baseline. This emphasizes that while the cell segmentation annotated by experts may be helpful in many ways, it should be used with caution as the gold standard for the cell segmentation task.

**Figure 5.**
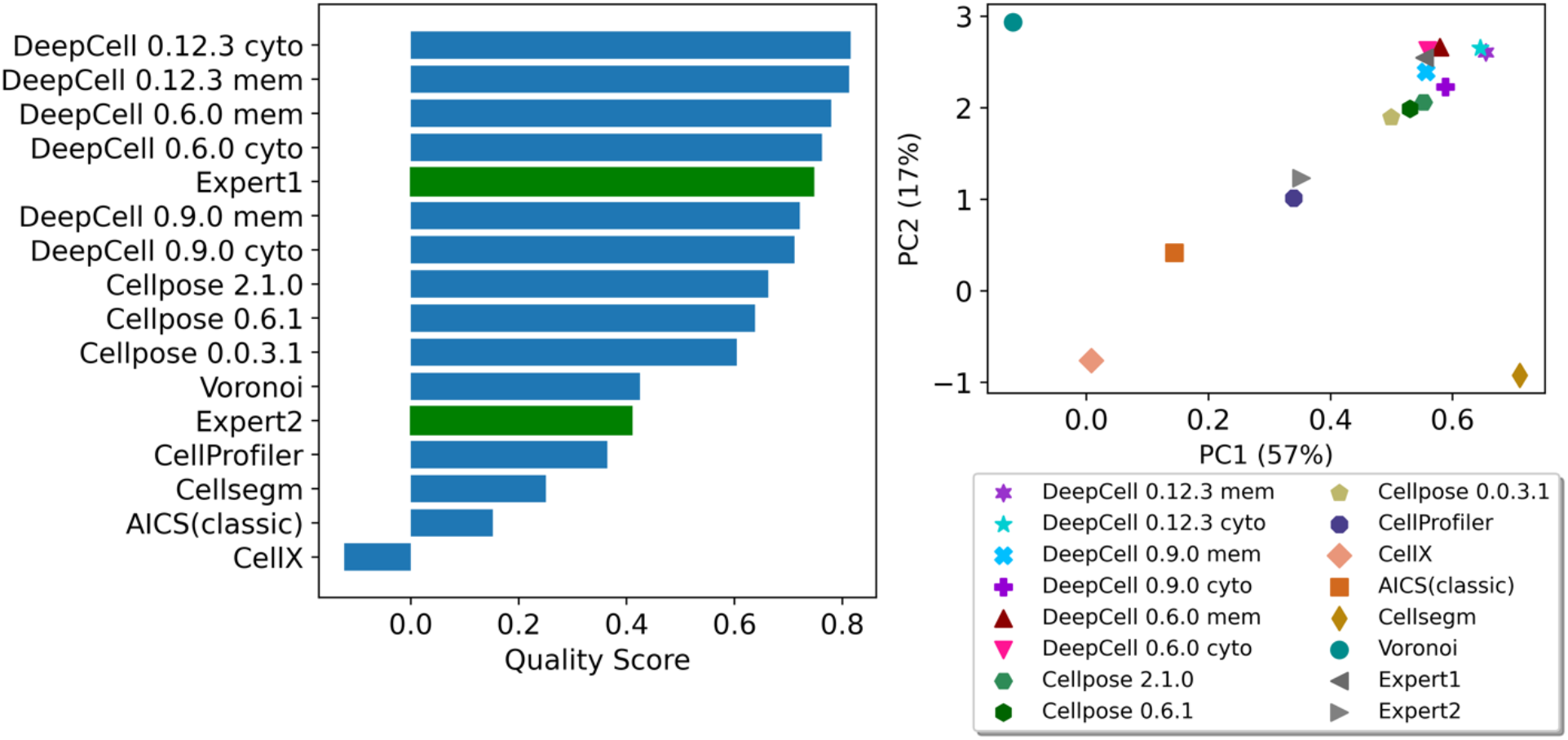
Quality score ranking and top 2 PCs of two expert-annotated segmentations comparing with results from all segmentation methods in Table 1. The accuracy of annotations from the Expert1 was reflected by our PCA model showing both high PC1 (good overall quality) and high PC2 (good tissue coverage) to the upper right corner among the group of deep learning-based model. The discrepancy between two experts was also captured showing inter-observer variability in the cell segmentation task.

We next sought to provide confidence that our quality scores obtained without a human expert would produce reliable results comparable to measures requiring human expert segmentation. To do this, we directly compared our quality scores with three benchmarks of cell segmentation quality (see Methods). These benchmarks were designed to be symmetric, meaning the results are the same by treating either human annotation or computer segmentation as the reference. The benchmarks were calculated for each method compared with each expert annotation in each image (Figure 6). We observed average Pearson correlation coefficients across the two images of 0.83, 0.86 and 0.72 between our quality score and the F1, PRAUC and SEG’ scores, respectively. Thus our quality scores are highly predictive of the benchmark scores that would have been obtained by comparison to expert annotations. Specifically, the deep learning-based methods on the upperright corner of three benchmarks show higher similarity with both expert annotations in either image. The non-deep learning-based methods, however, show lower agreement between quality score and either benchmark. This is because while these benchmarks mainly measure the similarity between annotation and segmentation from a single perspective (i.e., overlapping), our quality score measure bad segmentation performance in various ways (e.g., missing cells, cells too small, unmatched cell and nuclear masks). We also observed F1 scores around 0.7, PRAUC around 0.3 and SEG’ around 0.35 when comparing the two expert annotations, which are all lower than most deep learning-based methods when compared to either expert annotation. This inter-observer variance in terms of segmentation quality again echoes the conclusion from Figure 5 and supports the value of our quality scores as a measure of cell segmentation accuracy.

**Figure 6.**
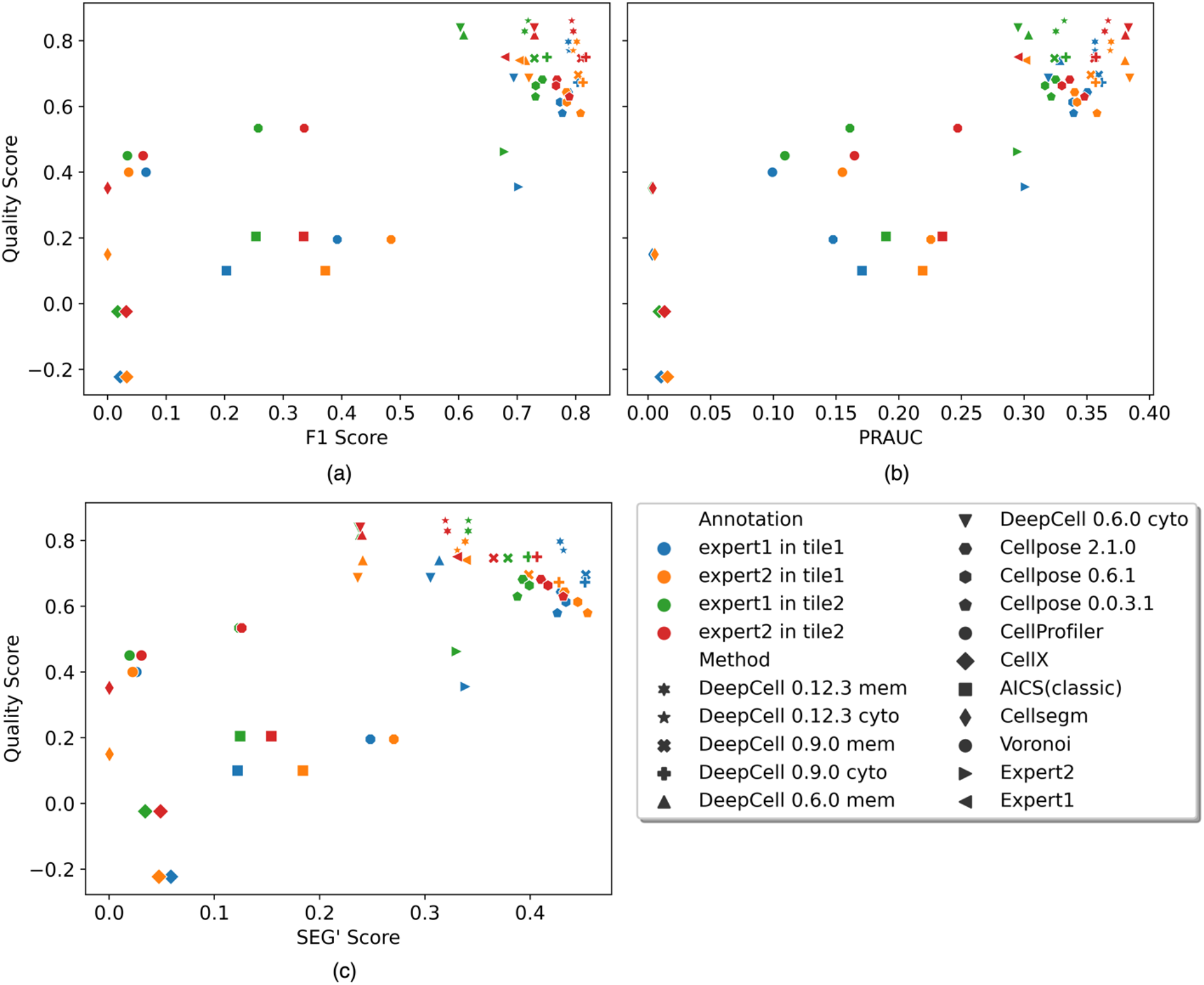
Comparing quality score with three benchmarks: F1 score (a), PRAUC (b) and SEG’ score (c). Each marker represents a segmentation or an expert annotation. Each color represents the benchmark was calculated with the marker’s segmentation/annotation against an expert annotation in a CODEX image (tile). Pearson correlation between our quality score and three benchmarks are 0.83, 0.86 and 0.72, respectively.

### Evaluating under-segmentation error

As shown in Figure 3, our quality scores appropriately reflect the degradation of cell segmentations quality that occurs after perturbing an input image. As a final test of the of the sensitivity of our quality scores, we directly degraded cell segmentations by two approaches. The first was to simulate under-segmentation, in which adjacent or overlapping objects that should be distinct are merged. It often occurs in cell and nucleus segmentation especially in dense tissues (Kromp et al., 2021). We simulated an under-segmentation scenario by merging cells in contact with each other. For this test, we started from masks generated by DeepCell 0.12.3, which is the best performer among all methods, on tiles from each of the CODEX datasets across all five tissue types, taking DAPI and E-cadherin as nuclear and cell membrane inputs (we selected one tile with enough cells to merge for each dataset). We sequentially merged pairs of contacting cells until the number of cells reached a target percent (either 90% or 60%) of the original cell number (note that cells that were merged were not allowed to merge again with another cell to better reflect typical undersegmentation performance, and that merging to 90% of the original cell number indicates 20% of cells in the original mask have been merged). For each pair of nuclei belonging to the merged cells, we applied morphological opening on the inverted mask to connect the gap between the nuclei to also simulate under-segmented nuclei (see Supplementary Figure 14).

As a second approach to degrading segmentation masks, we shifted the masks produced by DeepCell 0.12.3 relative to the original image. We shifted them by 0.1%, 1% and 50% of the average of the two dimensions of the image (for a 1000×1000 pixel image, shifting 0.1% is 1 pixel right and 1 pixel down). Presumably, results for 0.1% shift should be lower but similar to the original performance while the scores of the ones shifted 50% should be among the lowest.

We then ran our evaluation pipeline using DeepCell 0.12.3 on these degraded masks and compared the results with quality scores of all methods on the original images. The results in Figure 7 show that the quality scores reflect the degree of degradation produced. The breakdown of metrics in Supplementary Figure 15 illustrates that our evaluation method is sensitive to under-segmentation, but the degree of sensitivity expected depends upon the extent of variation in channel intensities among different cells (see Discussion). Notably, our evaluation method captured the degradation from merely shifting 0.1% with slightly lower quality score than the original.

**Figure 7.**
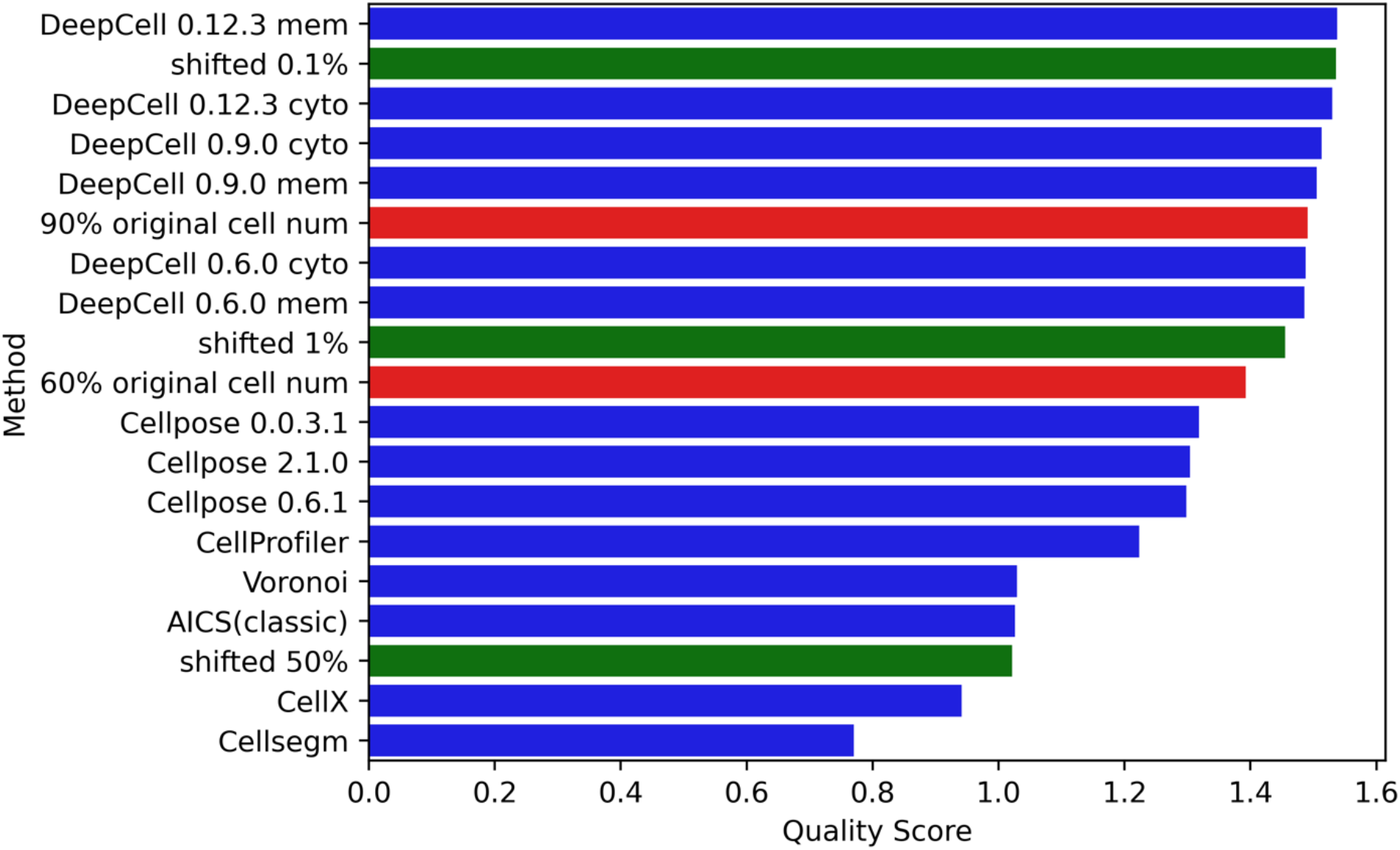
Quality score ranking of simulated masks with a range of under-segmentation error and shifting. Undersegmented masks (red) and shifted masks (green) were created from masks produced by DeepCell 0.12.3 using a cell membrane marker as described in the Methods.

### Evaluating similarity between results for different segmentation methods

Besides scoring each segmentation method separately, we also directly compared the segmentations produced by each pair of methods. We used two approaches: calculating a normalized distance between the single method metric vectors of two methods, and calculating a set of metrics from direct comparison of the segmentation masks from two methods (see Supplementary Methods). In each approach, we concatenated the metric matrices across all images from multiple data modalities for all pairs of methods and applied PCA to obtain PC1 values as a difference score. These scores were normalized by subtracting the PC1 value of a method compared with itself (i.e., the lowest possible difference metric value). Figure 8 shows heatmaps of the difference between the methods for the two sets of metrics. Higher values indicate more difference (lower similarity) between a pair of methods. Using the single method metrics (Figure 8a), all the deep learning-based models (DeepCell and Cellpose) are relatively close to each other, consistent with the results shown above. The results for DeepCell with the membrane and cytoplasm markers are also close. However, pairwise comparisons (Figure 8b) of the masks show larger differences in the results for DeepCell and Cellpose (including between the three versions of DeepCell), and somewhat larger differences between the results for membrane and cytoplasm markers.

**Figure 8.**
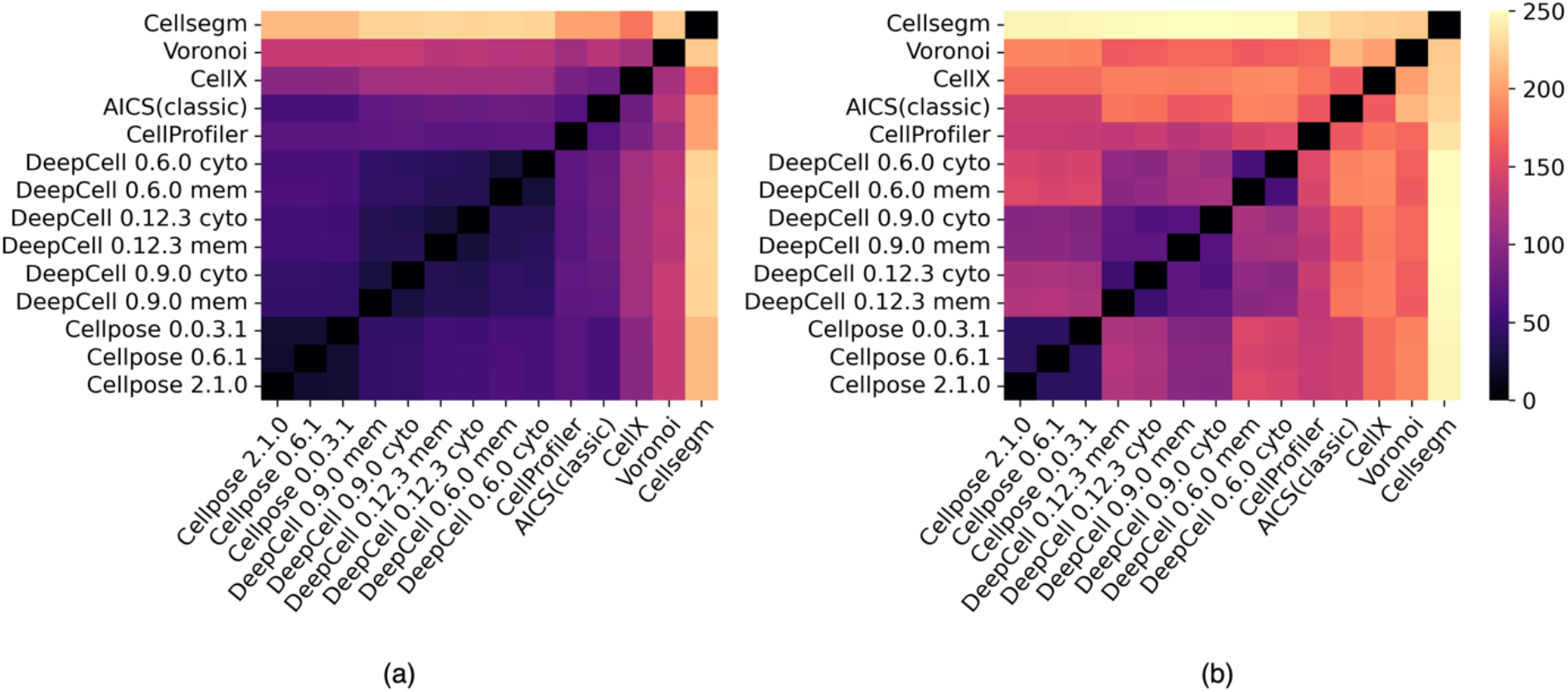
Heatmaps of difference scores between segmentation methods. A score of zero represents no difference (i.e. identical segmentations). Larger (lighter) values indicate greater differences between methods. (a) Heatmap generated by the difference between metrics of two single methods. (b) Heatmap generated by the pairwise metrics that directly compare two methods’ segmentations. The methods are ordered by clustering.

## Discussion

Many cell segmentation methods have been described but approaches to evaluate them have been limited. We have described here our design of a set of reference-free metrics to comprehensively measure the performance of segmentation methods. The metrics were developed based on a series of assumptions about the desirable characteristics of cell segmentation, especially for multichannel images. We also trained a PCA model using these metrics to compare all pre-trained segmentation models that we were able to identify. The results demonstrate the power of deep learning-based methods, with DeepCell and Cellpose performing the best. We also show that our metrics reflect the poorer performance expected for various image degradations, such as reducing pixel resolution, adding noise, and artificially increasing under-segmentation. Our evaluation approach also can be used to measure differences in the quality of different expert annotations. Our segmentation quality score highly correlates with three quality benchmarks that use expert annotations. Our open source tool has been incorporated into the image analysis pipeline for the HuBMAP project (HuBMAPConsortium, 2019).

The metrics are applicable across a range of image resolutions; the pixel size in the images we have used for evaluation range from 0.325 to 1 micron. Of course, the metrics (and the cell segmentation methods themselves) require on image resolution being sufficient to adequately resolve single cells.

An important consideration to note is that our metrics are based in large part on the assumption that tissue images used for evaluation contain different cell types that vary in their expression of different markers. If a tissue image is largely uniform, then our measures of segmented cell homogeneity will yield similar results regardless of the accuracy of cell segmentation masks. The greater the number of channels measured and the larger the differences between cell types, the more discriminating our metrics will be. The metrics also do not make any assumptions about allowable cell shapes. Measuring cell shape is an inherent problem for 2D tissue images where 3D cells are only seen in a thin layer in the z-axis. Cells belonging to the same cell type might be captured at different angles or heights and therefore display different shapes. Although our metrics of homogeneity at the cell level solve this issue indirectly (since cells from the same cell type but at different views should have similar channel compositions), segmentation methods that produce cells and nuclei with unexpected shapes are not directly penalized. Therefore, we plan in future work to incorporate metrics using spherical harmonic transform-based shape descriptors that have been shown to provide the best representation of cell and nuclear shapes (Ruan & Murphy, 2019). We also note that while this study was inspired by the multi-channel tissue images of the HuBMAP project, our metrics are also applicable to evaluate segmentation methods on cultured cells. However, as with tissue images, the sensitivity of the metrics depends on the number of channels and the diversity among individual cells.

To make our approach widely available, we provide an open source pipeline for calculating the segmentation metrics for a given set of images and a given segmentation method. Our evaluation pipeline provides a platform for users to choose segmentation methods for an individual image, tissue, and/or imaging modalities. While prior algorithmic approaches may have claimed the highest accuracy against different manually annotated training datasets, our method directly benchmarks them under the same set of measures.

The use of three-dimensional multi-channel tissue imaging is gradually growing. Some of the segmentation methods we have evaluated are capable of both 2D and 3D segmentation (see Methods), whereas algorithms such as the deep learning version of the Allen cell structure segmenter (Chen et al., 2020) and nnUNet (Isensee et al., 2021) only focus on segmenting 3D cell images. We are in the process of extending our evaluation work to 3D segmentation.

In closing, we note that in the future our quality scores could be included in the loss functions for training cell segmentation models in order to further improve model performance. They also provide a useful measure of image quality, since, within a given tissue and modality, better quality images can be expected to provide better segmentation results. This could potentially allow tissue images to be screened before acceptance by large projects such as HuBMAP.

## Acknowledgments

This work was supported in part by a grant from the National Institutes of Health Common Fund, OT2 OD026682 and by a traineeship to HC under training grant T32 EB009403.

