## Supplementary Information for "Evaluation of cell segmentation methods without reference segmentations"

Version 4, November 27, 2022

#### Supplementary Methods

##### Evaluation metrics not requiring reference segmentation

**Coverage Metrics**

The general assumption of coverage metrics is that a tissue image should be mostly occupied by cells except for readily detectable areas corresponding to ducts, blood vessels, or tissue borders. Therefore, we assume that a good segmentation should generate a mask that covers most areas of the tissue in an image.

*Number of cells per 100 square microns (NC)*

The principle is that cell density is a measure of the quality of both the image and the segmentation method. The pixel size in square microns varies in different image modalities. To calculate the density, pixel sizes in the X and Y dimensions are directly obtained from OME-TIFF metadata. The metric is defined as

$$\mathrm{NC} = \frac{n_c}{s_p \times n_p} \times 100$$

where $n_c$ is the number of cells in the cell mask, $s_p$ is the size of one pixel in squared microns, $n_p$ is the total number of pixels in the corresponding image.

Given the binary foreground-background separation (see **Foreground-background separation**) of a tissue image, we designed the following three coverage metrics with the general assumption that an accurate segmentation mask will cover most tissue areas (i.e., the foreground) but few background areas.

*Fraction of image foreground occupied by cells (FFC).*

$$\mathrm{FCF} = \frac{a_{cf}}{a_f}$$

where $a_{cf}$ is the area of a cell mask in the foreground and $a_f$ is the area of the foreground.

*Fraction of image background occupied by cells (FBC)*

$$\mathrm{FBC} = \frac{a_{cb}}{a_b}$$

where $a_{cb}$ is the area of a cell mask in the background and $a_f$ is the area of the background.

1 minus FBC is used as the metric, so that larger values indicate better segmentation.

*Fraction of cell mask in foreground (FCF)*

$$\mathrm{FCF} = \frac{a_{cf}}{a_c}$$

where $a_{cf}$ is the area of a cell mask in the foreground and $a_c$ is the area of the cell mask.

After matching the cell and nuclear masks (see **Mask processing**), we calculated the fraction of matched cells and nuclei, assuming that segmented cells and nuclei should have a one-to-one correspondence relationship.

*Fraction of match between cells and nuclei (FMCN)*

$$\text{FMCN} = \frac{n_m}{n_{cmi} + n_{nmi} + n_m}$$

where $n_m$ is the number of matched cells and nuclei, $n_{cmi}$ is the number of mismatched cells, $n_{nmi}$ is the number of mismatched nuclei.

**Homogeneity Metrics**

To measure the homogeneities of pixel intensities and integrated cell intensities from a segmentation method, we developed another category called homogeneity metrics. At the pixel level, the principle is that the pixels outside of the cells but still within the image foreground should be similar in protein composition (i.e., should consist of extracellular matrix of similar composition). The coefficient of variation (CV) of the foreground pixels outside the cells (Supplementary Figure 16a) for each channel was calculated and then the average CV across all channels was taken.

*Average CV of foreground pixels outside the cells (ACVF).*

$$\text{ACVF} = \frac{1}{n_h}\sum_{i=1}^{n_h}\frac{\sigma_{if}}{\mu_i}$$

where $n_h$ is the total number of image channels, $\sigma_{if}$ is the standard deviation of pixel intensities of the $i^{th}$ channel in the foreground outside the cells, $\mu_i$ is the mean of all pixel intensities in the $i^{th}$channel.

The reciprocal of ACVF+1 is used as the metric, so that higher values indicate better segmentation and its upper bound is 1.

We applied the Principal Component Analysis (PCA) on the matrix of all foreground outside-the-cell pixel intensities across all channels after z-score standardization on each channel was applied and the fraction of variance explained by the first principal component was calculated. A higher value of the fraction stands for a more conserved relationship across channels in the foreground outside the cell areas (Supplementary Figure 16a).

*Fraction of first PC of foreground pixels outside the cells (FPCF).*

$$\mathrm{FPCF} = \frac{\lambda_{f1}}{tr(\Sigma_f)}$$

where $\lambda_{f1}$ is the variance explained by the first principle component (PC) across channels for the foreground pixels outside the cells, $\Sigma_f$ is the covariance matrices from PCA analysis on the foreground pixels outside the cells across all channels, $tr(\quad)$ is trace calculation on a matrix.

At the cell level, the principles are that (a) cell types should be roughly similar in composition across all channels, (b) cell types can be approximated by cell clusters, (c) the average CV across all clusters for a given number of clusters is a proxy for similarity in composition within cell types, and (d) averaging the average CV across all clusters across different numbers of the cluster is also a proxy for similarity in composition within cell types. Cell types were defined using KMeans clustering performed on the mean cell intensities across all channels after applying z-score standardization for each channel. Since the number of cell types in an arbitrary image is unknown, K, the number of clusters, was varied from 1 to 10. Note that the KMeans clustering was applied on the cell mask to define the cell type, which generated cell type labels of each number of clusters used in the following metric calculation for each cellular component mask (Supplementary Figure 16b).

For the clusters from each K value, the average CV for each cluster across channels was calculated followed by an average weighted by the cluster size. The final metric was calculated as the average of the weighted average coefficient of variation across all clusters over 1 to 10 clusters.

*Average of weighted average CV of cell type intensities over 1-10 clusters (ACVC)*

$$\text{ACVC} = \frac{1}{K_{max} - K_{min} + 1} \sum_{K=K_{min}}^{K_{max}} \left( \frac{1}{n_c} \sum_{k=1}^{K} (CV)_k \cdot n_k \right)$$

$$(CV)_k = \frac{1}{n_h} \sum_{i=1}^{n_h} \frac{\sigma_{ki}}{\mu_{ki}}$$

where $K_{min}$ is the smallest number of clusters, $K_{max}$ is the largest number of clusters, $K$ is the current number of clusters, $n_c$is the total number of cells, $CV_k$ is the coefficient of variation of mean cell intensities in $k^{th}$ cluster, $\sigma_{ki}$ and $\mu_{ki}$ is the standard deviation and the mean of mean cell intensities of $k^{th}$ cluster in $i^{th}$ channel respectively, $n_k$ is the number of cells in $k^{th}$ cluster, $n_h$ is the total number of image channels.

The reciprocal of ACVC+1 is used as the metric, so that higher values indicate better segmentation and its upper bound is 1.

A similar measure was derived using principal component analysis. The first two principles are the same as above but (c) the fraction of variance accounted for by the first principal component of the cells in each cluster is a proxy for similarity in composition of each cell type, and (d) averaging this fraction over different numbers of clusters is also a proxy for similarity of each cell type. We, therefore, applied PCA on the mean cell intensities across z-score standardized channels of each number of cluster and calculated the average fraction of variance accounted for by the first principal component across all numbers of clusters. The output vector was averaged over different K values to get the final metric.

*Average of weighted average fraction of the first PC of cell type intensities over 1-10 clusters*

$$\text{FPCC} = \frac{1}{K_{max} - K_{min} + 1} \sum_{K=K_{min}}^{K_{max}} \left( \frac{1}{n_c} \sum_{k=1}^{K} \frac{\lambda_{k1}}{tr(\Sigma_k)} \cdot n_k \right)$$

where $K_{min}$ is the smallest number of clusters, $K_{max}$ is the largest number of clusters, $K$ is the current number of clusters of choice, $n_c$ is the total number of cells, $\lambda_{k1}$ is the variance explained by the first principle component in $k^{th}$ cluster, $\Sigma_k$ the covariance matrices from PCA on mean cell intensities across all channels in $k^{th}$ cluster, $n_k$ is the number of cells in the current cluster.

We also incorporated the Silhouette score as a metric to indicate the similarity of composition across cell types. Since the Silhouette score equals 1 when there is only one cluster, we started with two clusters to calculate this metric.

*Average of Silhouette score of clustering over 2-10 clusters (AS)*

$$\text{AS} = \frac{1}{K_{max} - K_{min} + 1} \sum_{K=K_{min}}^{K_{max}} \left( \frac{1}{n_c} \sum_{i=1}^{n_c} \frac{b(i) - a(i)}{\max(a(i), b(i))} \right)$$

$$\text{a}(i) = \frac{1}{|K_i| - 1} \sum_{j \in K_i, i \neq j} d(i,j)$$

$$\text{b}(i) = \min_{l \neq i} \frac{1}{|K_l|} \sum_{j \in K_l} d(i,j)$$

where $K_{min}$ is the smallest number of clusters, $K_{max}$ is the largest number of clusters, $K$ is the current number of clusters of choice, $a(i)$ is the average distance between cell $i$ and all the other cells $j$ in the cluster $K_i$ to which cell $i$ belongs, $b(i)$ is the minimum average distance from cell $i$ to all cells $j$ in all clusters $K_l$ to which $i$ does not belong, $n_c$ is the total number of cells.

In addition, we assumed that properly segmented images should have similarly sized cells.

*Standard Deviation of cell size (CSSD)*

$$\text{CSSD} = \sqrt{\frac{\sum_{i=1}^{n_c}(s_i - \mu_s)^2}{n_c}}$$

where $s_i$ is the size of $i^{th}$ cell, $\mu_s$ is the mean size of all cells, $n_c$ is the total number of cells.

The reciprocal of ln(CSSD)+1 is used as the metric, so that higher values indicate better segmentation and its upper bound is 1. The natural logarithm is taken to close the large difference of CSSD between methods.

#### Foreground-background separation

To assess the coverage of the segmentation mask, each image was separated into regions consisting primarily of foreground and background. To do this, mean thresholding was applied to the nuclear, cytoplasmic, and cell membrane channels respectively, followed by performing two rounds of morphological closing with disk kernel of radius 1 and 10, respectively, on the union of three thresholded binary images. Each round consists of a foreground closing to bridge the small (first round) or large (second round) gaps between cells within the tissue and a background closing on the inverted image to reunite the overly scattered background. An area closing was subsequently applied and considered those areas with 0 values as the extra-cellular matrices in the foreground (and assign 1) if they are less than 5000 pixels, which can be tuned according to the image resolution. To further correct the resulting rounded boundaries, a morphological geodesic active contour was applied on the inverted image with current background areas as seeds, followed by an area closing to remove the small dots in the background. The final output is a binary mask with 1 as foreground and 0 as background (Supplementary Figure 17).

#### Evaluation metrics for the similarity between two segmentation methods

**Difference between the evaluation metrics of two masks**

We assumed that the values of evaluation metrics for a single segmentation should be close for two similar segmentation methods. We took the absolute difference of each pair of metrics between two masks as the first set of metrics to evaluate the similarity between the two methods.

**Pairwise metrics**

We designed and adopted metrics for comparing two segmentation methods. Some of the following pairwise metrics were summarized by (Taha & Hanbury, 2015).

**Number of Matched Cells (NMC)**

We hypothesized that a large number of segmented cells should be overlapped for similar segmentation methods. We defined the metrics as the number of cell pairs from two segmentations that have an overlap fraction greater than a fraction of the larger cell in each pair. When one cell in a mask overlaps with multiple cells in another mask, only the one with the largest overlap is considered matched.

$$\mathrm{NMC}(S_1, S_2) = \sum_i \sum_j \mathrm{I}_{|x_i \cap y_j| > p \max(|x_i|, |y_j|)}$$

where $S_1$, $S_2$ are two segmentation masks, $\mathrm{I}$ is the indicator function, $x_i$ and $y_j$ is $i^{th}$ and $j^{th}$ cell in $S_1$ and $S_2$ respectively, overlap fraction threshold $p$ is a free parameter. We took $p = 0.5$.

**Threshold AUC (TAUC)**

We assumed that similar segmentation methods should have masks with an overall good overlap of cells for varying overlap fraction thresholds. We calculated the area under the curve of the percentage of overlapping cells vs overlap fraction threshold while the threshold ranged from 0 to 1. The percentage of overlapping cells is the number of cell pairs that have an overlap fraction greater than the threshold, normalized by the smaller cell numbers of two masks.

**Average Jaccard index (AJI)**

We assumed that the overall similarity between matched cells should be high for similar segmentation methods. An average Jaccard index (Jaccard, 1912; Tanimoto, 1958) over all pairs of matched cells is thus calculated as a metric.

$$AJI(S_1, S_2) = \frac{1}{n_m} \sum_i \frac{|x_i \cap y_i|}{|x_i \cup y_i|}$$

where $S_1$, $S_2$ are two segmentation masks, $x_i$ and $y_i$ is $i^{th}$ pair of matched cells, $x_i \in S_1$, $y_i \in S_2$, $n_m$ is the total number of matched pairs.

**Average Dice Coefficient (ADC)**

With the same assumption as for the Jaccard index, we also calculated an average Dice coefficient (Dice, 1945; Sorensen, 1948) over all pairs of matched cells.

$$ADC(S_1, S_2) = \frac{1}{n_m} \sum_i \frac{2|x_i \cap y_i|}{|x_i| + |y_i|}$$

where $S_1$, $S_2$ are two segmentation masks, $x_i$ and $y_i$ is $i^{th}$ pair of matched cells, $x_i \in S_1$, $y_i \in S_2$, $n_m$ is the total number of matched pairs.

**Average Hausdorff Distance (AHD)**

We assumed that each pair of matched cells should have a close spatial distance for similar segmentation methods. We selected Hausdorff distance (Birsan & Tiba, 2005; Rockafellar & Wets, 2009) as the metric to measure the spatial distance between cells.

$$AHD(S_1, S_2) = \frac{1}{n_m} \sum_i \max\left( \sup_{a \in x_i} \inf_{b \in y_i} d\,(a, b), \sup_{b \in y_i} \inf_{a \in x_i} d\,(a, b) \right)$$

where $S_1$, $S_2$ are two segmentation masks, $x_i$ and $y_i$ is $i^{th}$ pair of matched cells, $x_i \in S_1$, $y_i \in S_2$, $a$, $b$ are the pixels in $x_i$ and $y_i$ cells respectively, $d$ is the distance between two pixels, and $n_m$ is the total number of matched pairs.

**Bidirectional Consistency Error (BCE)**

We assumed two similar segmentation methods should be consistent with the regions of each pixel that belongs. Bidirectional Consistency Error (Martin, 2002) measures the error between two segmentation methods in terms of the regions over all pixels.

$$BCE(S_1, S_2) = \frac{1}{n_p} \max\left(\sum_i E(S_1, S_2, p_i), \sum_i E(S_2, S_1, p_i)\right)$$

$$E(S_1, S_2, p_i) = \frac{|R(S_1, p_i) - R(S_2, p_i)|}{R(S_1, p_i)}$$

where $S_1$, $S_2$ are two segmentation masks, $p_i$ is $i^{th}$ pixel in the image, $R(S_j, p_i)$ is the area of the region in segmentation $S_j$ that contains pixel $p_i$, $n_p$ is the total number of pixels in the image.

**Error Matrix**

Considering one mask as the reference and the other as the query, we have:

- True Positive (TP): Total number of overlapping pixels in matched cells
- False Positive (FP): Total number of pixels that are in query mask but not in reference mask
- True Negative (TN): Total number of all pixels outside both masks
- False Negative (FN): Total number of pixels that are in reference mask but not in query mask

The following metrics using the error matrix to calculate are symmetric no matter which mask is considered as the reference.

**Cohen's Kappa (CK)**

Cohen's Kappa (McHugh, 2012) measures the agreement between raters. In this study, each segmentation method is a rater to the image. We assumed that two segmentation methods should agree with each other in terms of Cohen's Kappa if they are similar.

$$CK(S_1, S_2) = \frac{f_a - f_c}{n_p - f_c}$$

$$f_a = TP + TN$$

$$f_c = \frac{(TN + FN)(TN + FP) + (FP + TP)(FN + TP)}{n_p}$$

where $S_1$, $S_2$ are two segmentation masks, $n_p$ is the total number of pixels in the image.

**Mutual Information (MI)**

We assumed that one segmentation should provide most of the information for the other when they are similar. We applied mutual information (Shannon, 1948) to measure the reduction in uncertainty of one segmentation to another.

$$MI(S_1, S_2) = H(S_1) + H(S_2) - H(S_1, S_2)$$

$$H(S) = -\sum_i p(S^i) \log p\left(S^i\right)$$

$$H(S_1, S_2) = -\sum_{ij} p(S_1^i, S_2^j) \log p\left(S_1^i, S_2^j\right)$$

$$p(S_1^1, S_2^1) = TP/n_p \quad p(S_1^1) = (TP + FN)/n_p$$

$$p(S_1^1, S_2^2) = FN/n_p \quad p(S_1^2) = (TN + FN)/n_p$$

$$p(S_2^1, S_1^2) = FP/n_p \quad p(S_2^1) = (TP + FP)/n_p$$

$$p(S_1^2, S_2^2) = TN/n_p \quad p(S_2^2) = (TN + FP)/n_p$$

where $S_1$, $S_2$ are two segmentation masks, $n_p$ is the total number of pixels in the image; superscript = 1 means segmented cell region in the mask, 2 means segmented background region in the mask.

**Variation of Information (VOI)**

We assumed that similar segmentation methods should provide nearly amount of information. We applied the variation of information (Arabie & Boorman, 1973; Meilă, 2003; Unruh & Zurek, 1989) to measure the amount of information lost (or gained) when changing from one segmentation to another.

$$VOI(S_1, S_2) = H(S_1) + H(S_2) - 2MI(S_1, S_2)$$

where $S_1$, $S_2$ are two segmentation masks, $MI$ is the mutual information metric.

**Volumetric Similarity (VS)**

We assumed that similar segmentation methods should generate segmentation masks with similar area. While Average Hausdorff Distance is majorly considering the similarity of boundaries and shapes between two cell masks, we also applied volumetric similarity (Taha & Hanbury, 2015) to measure the similarity in terms of segmented area.

$$VS(S_1, S_2) = 1 - \frac{|FN - FP|}{2 * TP + FP + FN}$$

where $S_1$, $S_2$ are two segmentation masks.

#### Supplementary Tables

Supplementary Table 1 Summary of nuclear, cytoplasmic, and cell membrane channels for each imaging modality

| | Number of images (tiles) | Nuclei | Cytoplasm | Cell Membrane |
|---|---|---|---|---|
| CODEX-Standford | 96 | Hoechst1 | Cytokeratin | CD45 |
| CODEX-Univ. of Florida | 297 | DAPI | CD107a | E-cadherin |
| MIBI | 10 | HH3 | PanKeratin | E-cadherin |
| Cell DIVE | 221 | DAPI | Cytokeratin | P-cadherin |
| IMC | 13 | lr191/Histone | SMA | HLA-ABC |

Supplementary Table 2 Summary of markers as cell segmentation input

| Marker | Target | Subcellular Localization | Source |
|---|---|---|---|
| DAPI | DNA | Nucleus | (Kapuscinski, 1995) |
| Hoechst | DNA | Nucleus | (Latt & Stetten, 1976; Latt et al., 1975) |
| Ir191 | DNA | Nucleus | (Sumatoh et al., 2017) |
| Histone/HH3 | histone | Nucleus | https://www.uniprot.org/uniprotkb/B4E380/entry#subcellular_location |
| E-cadherin | CDH1 | Cell membrane | https://www.uniprot.org/uniprotkb/P12830/entry#subcellular_location |
| HLA-ABC | MHC class I antigen | Cell membrane | https://www.uniprot.org/uniprotkb/O19689/entry#subcellular_location |
| P-cadherin | CDH3 | Cell membrane | https://www.uniprot.org/uniprotkb/P22223/entry#subcellular_location |
| CD45 | Receptor-type tyrosine-protein | Cell membrane | https://www.uniprot.org/uniprotkb/P08575/entry#subcellular_location |
| SMA | Smooth muscle actin | Cytoplasm | https://www.uniprot.org/uniprotkb/P63267/entry#subcellular_location |
| CD107a | LAMP1 | Cytoplasm | https://www.uniprot.org/uniprotkb/P11279/entry#subcellular_location |
| PanKeratin/ Cytokeratin | intermediate filament | Cytoplasm | https://www.uniprot.org/uniprotkb/Q16195/entry#subcellular_location |

### Supplementary Figures

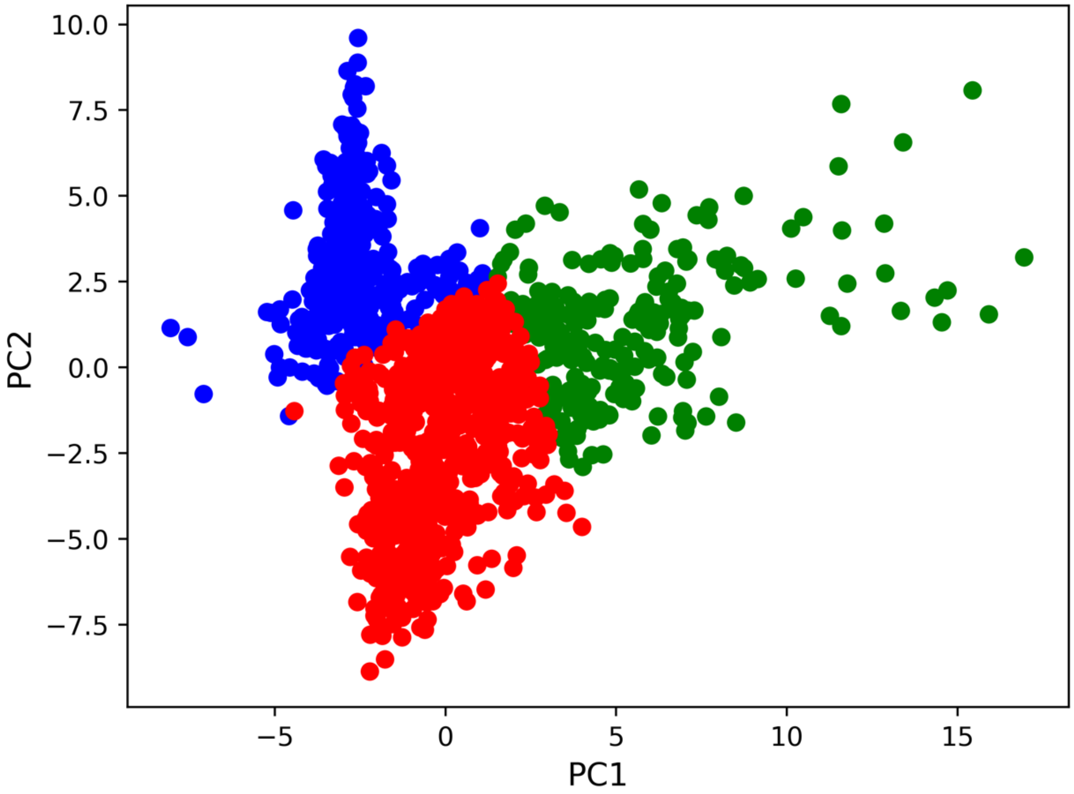


Supplementary Figure 1 To select grided tiles with dense cells, we calculated three image quality metrics for every channel of each tile of Cell DIVE grided images: 1. Signal To Noise Otsu: The Otsu method is used to choose an intensity threshold for each channel and the ratio of the average intensities above and below the threshold is calculated for each channel. 2. Signal to Noise Z-Score: The ratio of the mean intensity to the standard deviation of intensity is calculated for each channel. 3. Total Intensity: The sum of intensities is calculated for each channel. With three metrics of every channel, we constructed a matrix with each row as a tile and each column as a metric of a channel of this tile. We applied KMeans clustering on this matrix with number of cluster=3. To better illustrate the clustering result, we applied PCA on the same matrix. Each dot on the figure above is a tile. The tiles in blue cluster are nearly empty. The red tiles have few cells or extra-cellular matrix. The green cluster contains 221 tiles with abundant cells and tissue areas, which we eventually used as Cell DIVE data in our segmentation-evaluation pipeline.

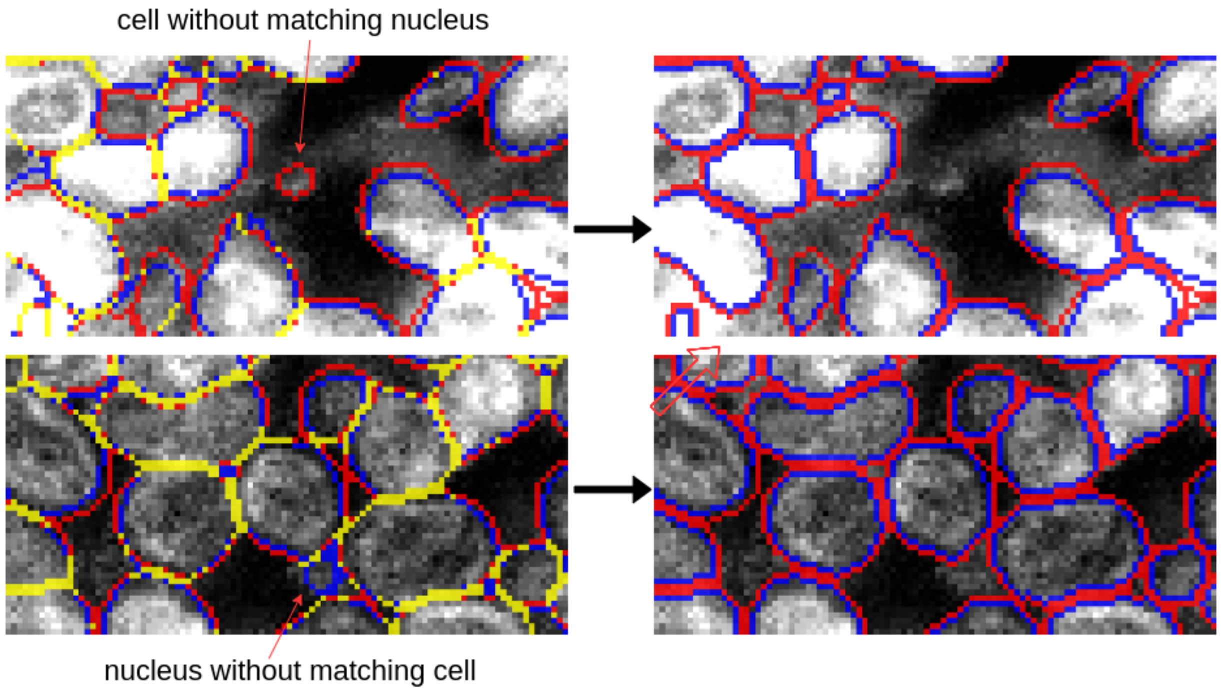


Supplementary Figure 2 The illustration of the postprocessing mask repair step. The red pixels are the cell boundaries from the cell segmentation. The blue pixels are the nuclear boundaries from the nuclear segmentation. Both segmentations are from the same method (DeepCell 0.9.0 cell membrane). The yellow pixels are the overlap between cell and nuclear boundaries before repair. In the repair step, we trimmed the nuclear boundaries outside and on the cell boundaries to make sure the nuclei are completely within the cells (no yellow pixels remain). Cell without matching nucleus and nucleus without matching cells were removed.

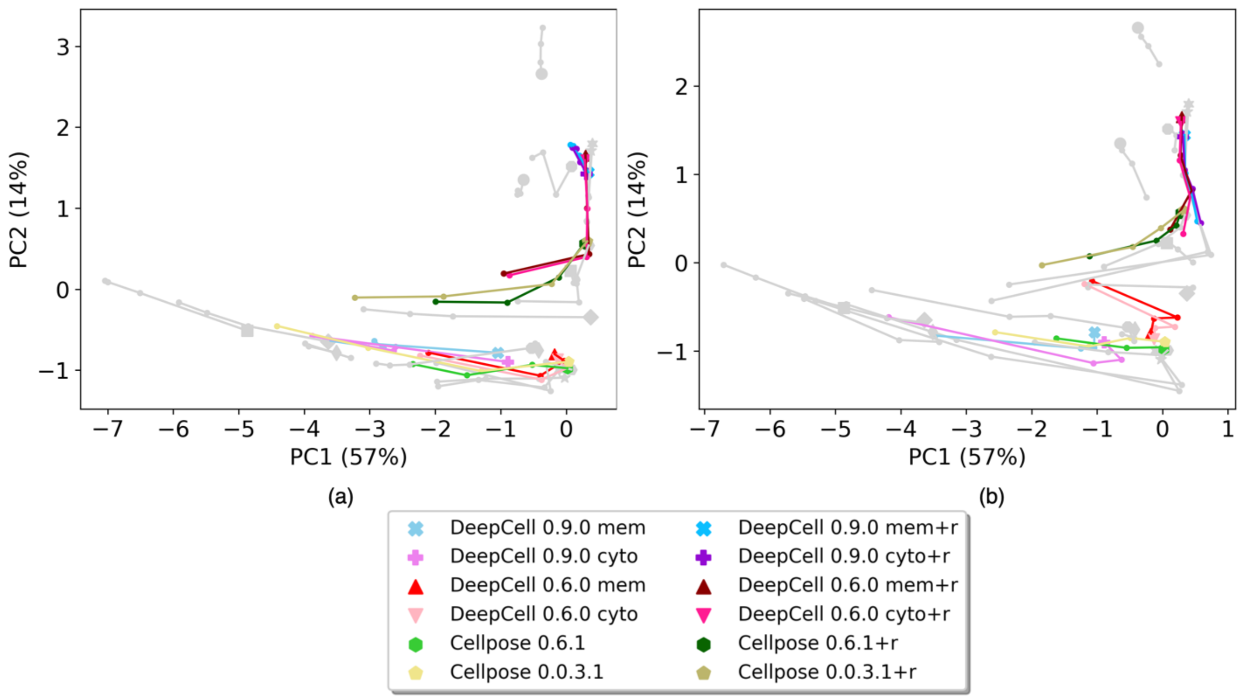


Supplementary Figure 4 Top 2 Principal components of previous versions of DeepCell and Cellpose on images from all modalities. Each method is represented by a unique color. All other methods are shown in gray. (a) Results from images with Gaussian noise perturbation. (b) Result from images with downsampling.

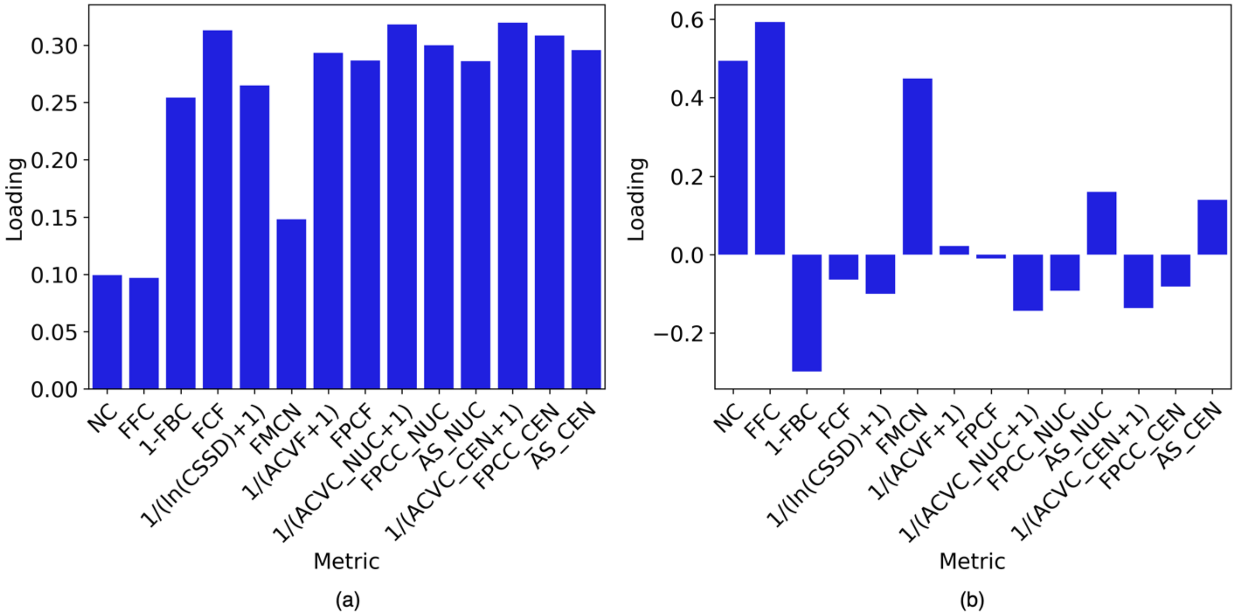


Supplementary Figure 4 Factor loadings of PCA model trained by all segmentation masks across all modalities from all methods. (a) Loadings of PC1 (b) Loadings of PC2

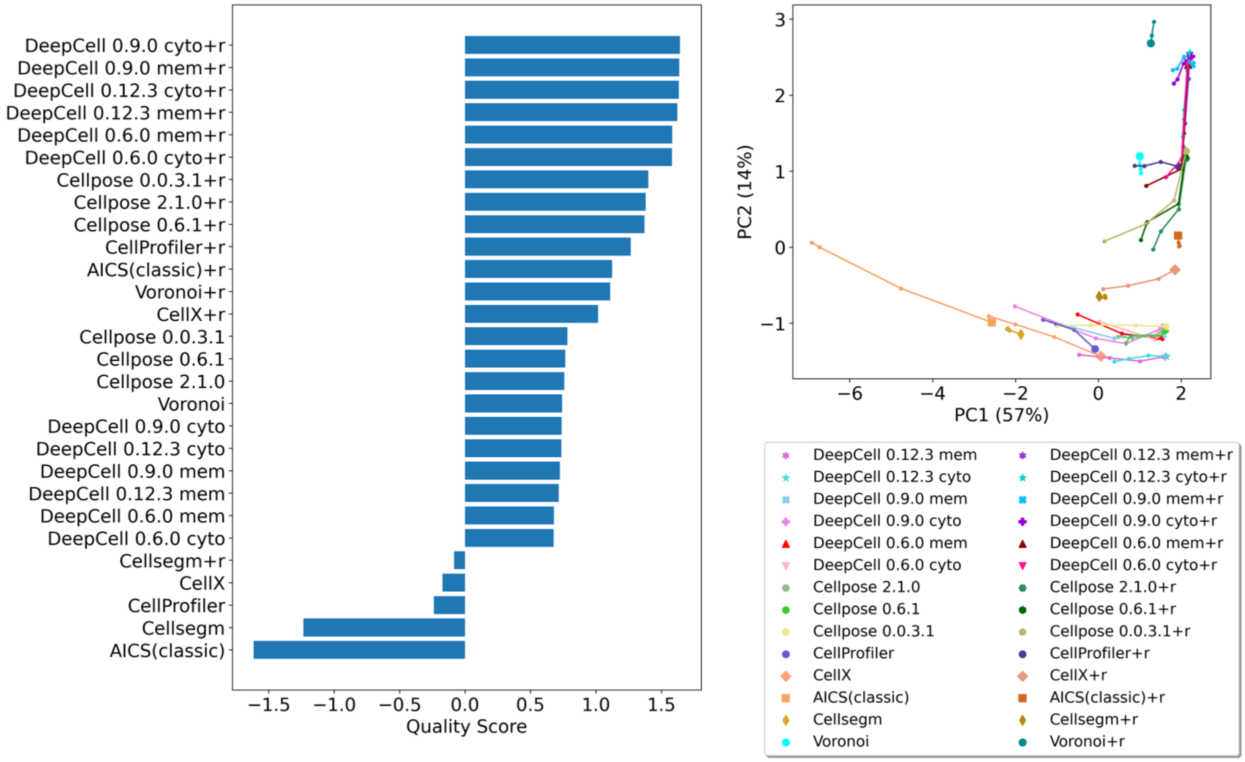


Supplementary Figure 5 Ranking of segmentation quality scores of all methods on CODEX images

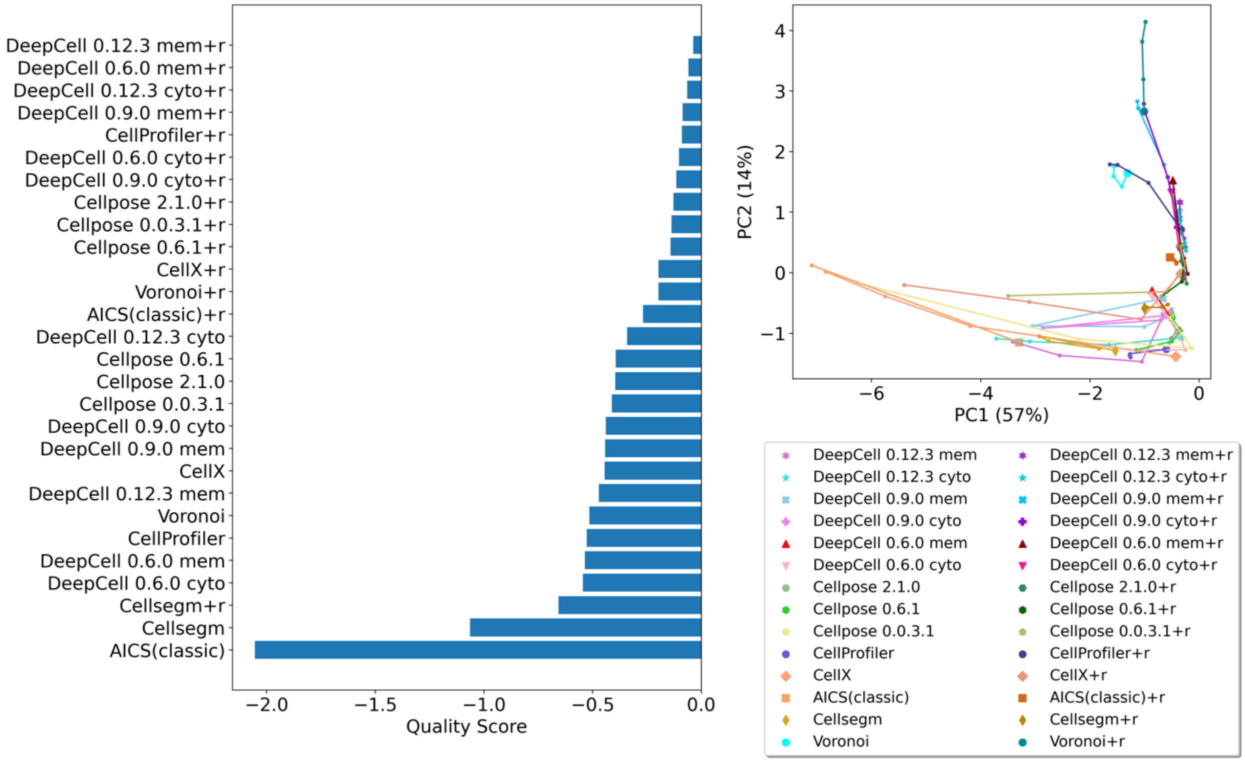


Supplementary Figure 6 Ranking of segmentation quality scores of all methods on Cell DIVE images

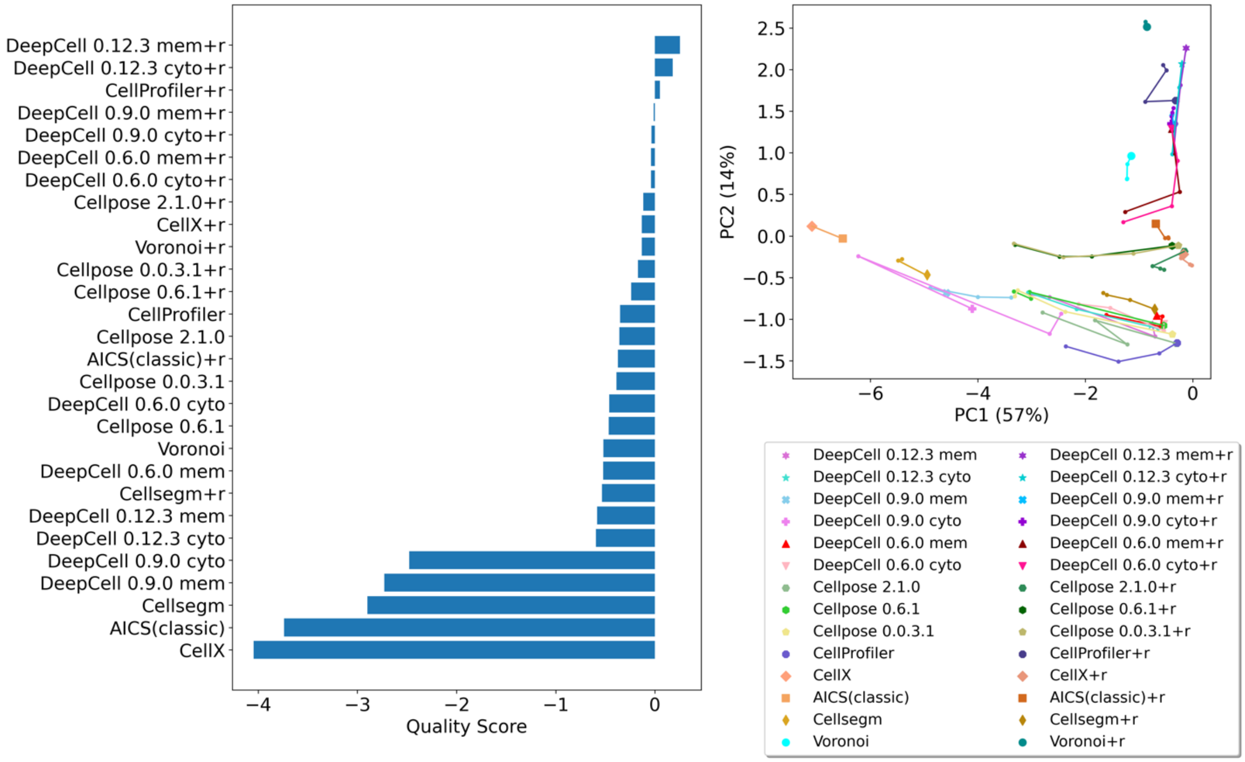


Supplementary Figure 7 Ranking of segmentation quality scores of all methods on IMC images

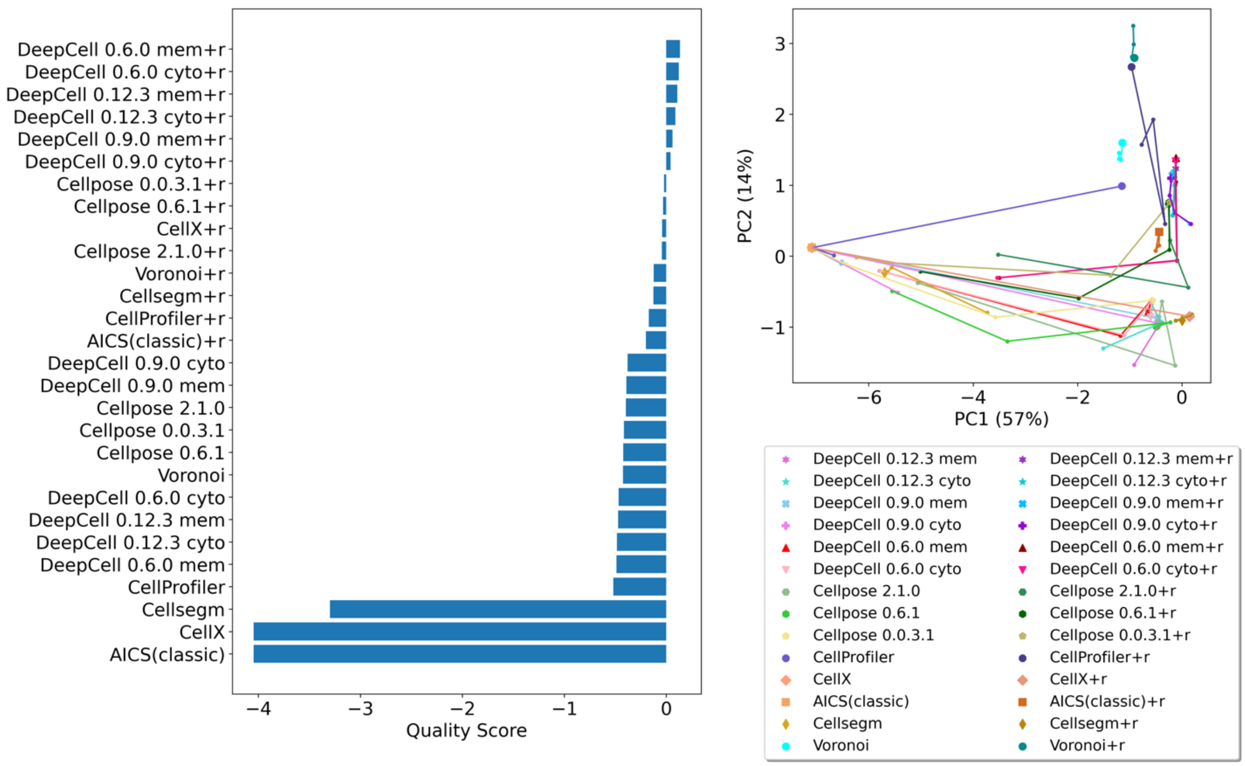


Supplementary Figure 8 Ranking of segmentation quality scores of all methods on MIBI images

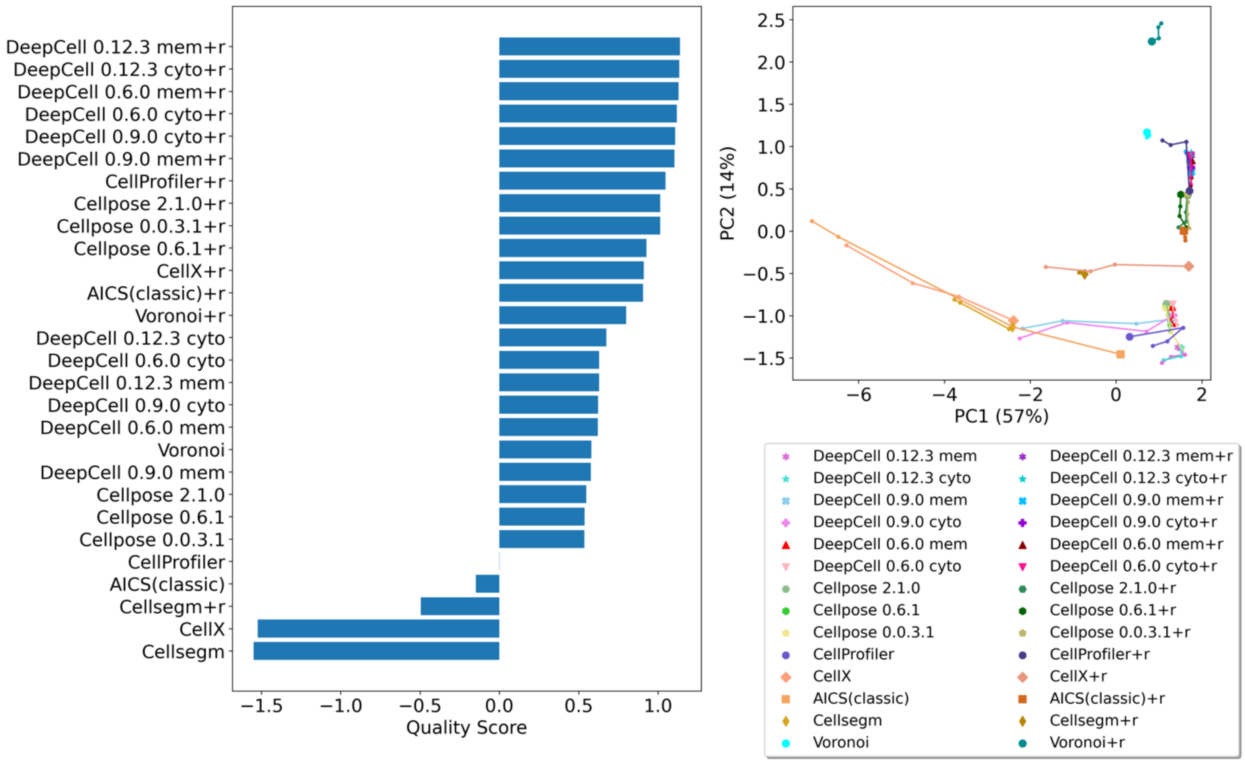


Supplementary Figure 9 Ranking of segmentation quality scores of all methods on Small Intestine images in CODEX datasets.

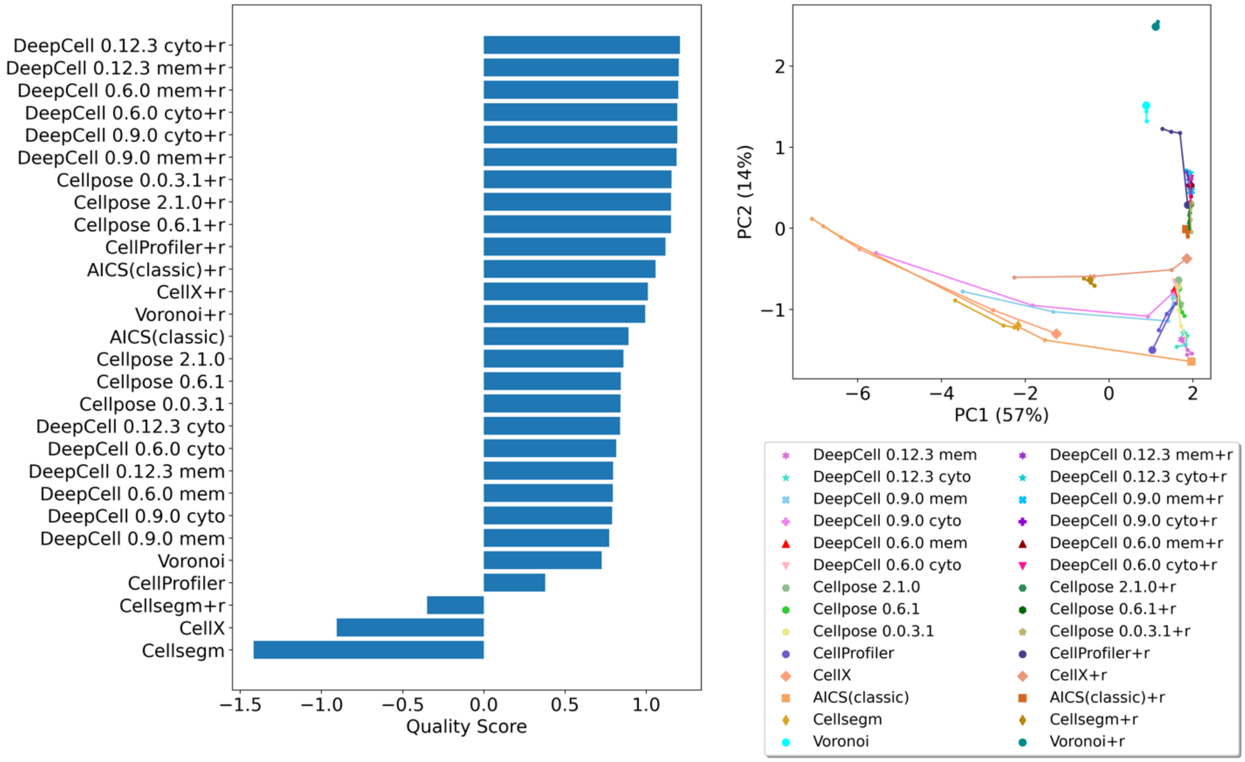


Supplementary Figure 10 Ranking of segmentation quality scores of all methods on Large Intestine images in CODEX datasets.

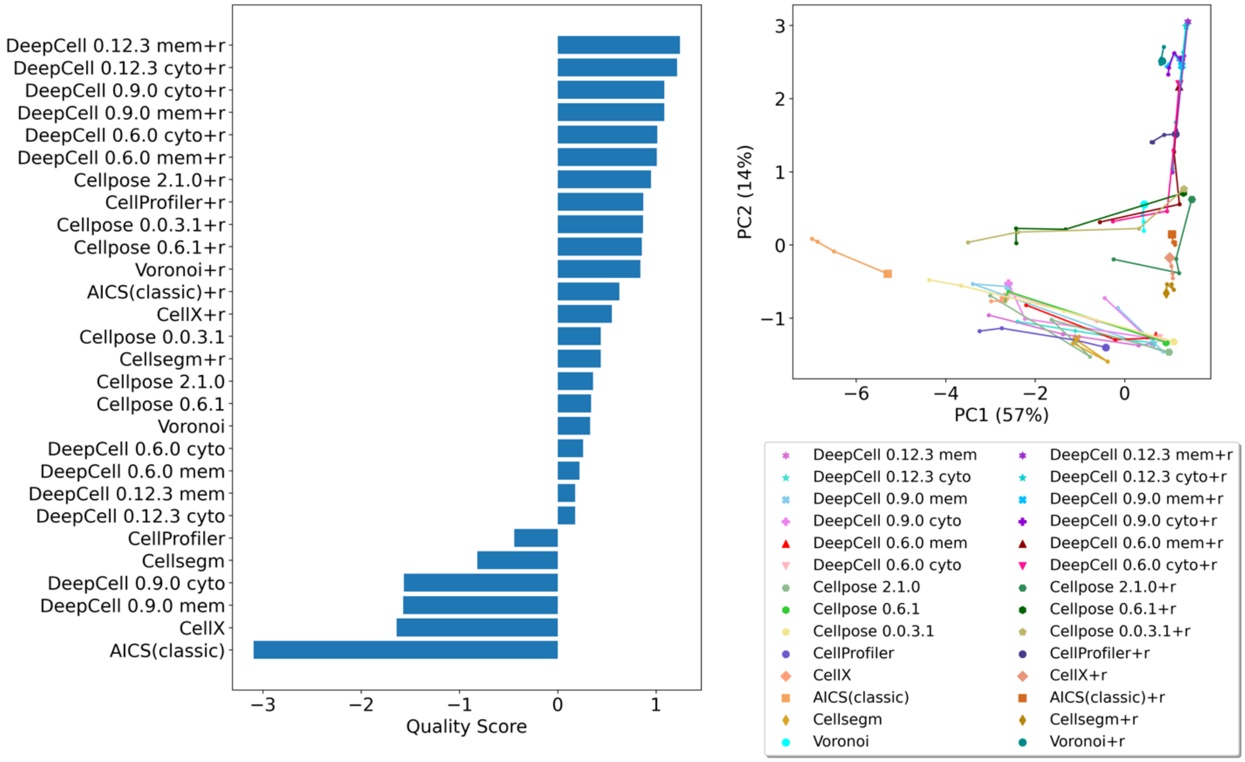


Supplementary Figure 11 Ranking of segmentation quality scores of all methods on Spleen images in CODEX and IMC datasets.

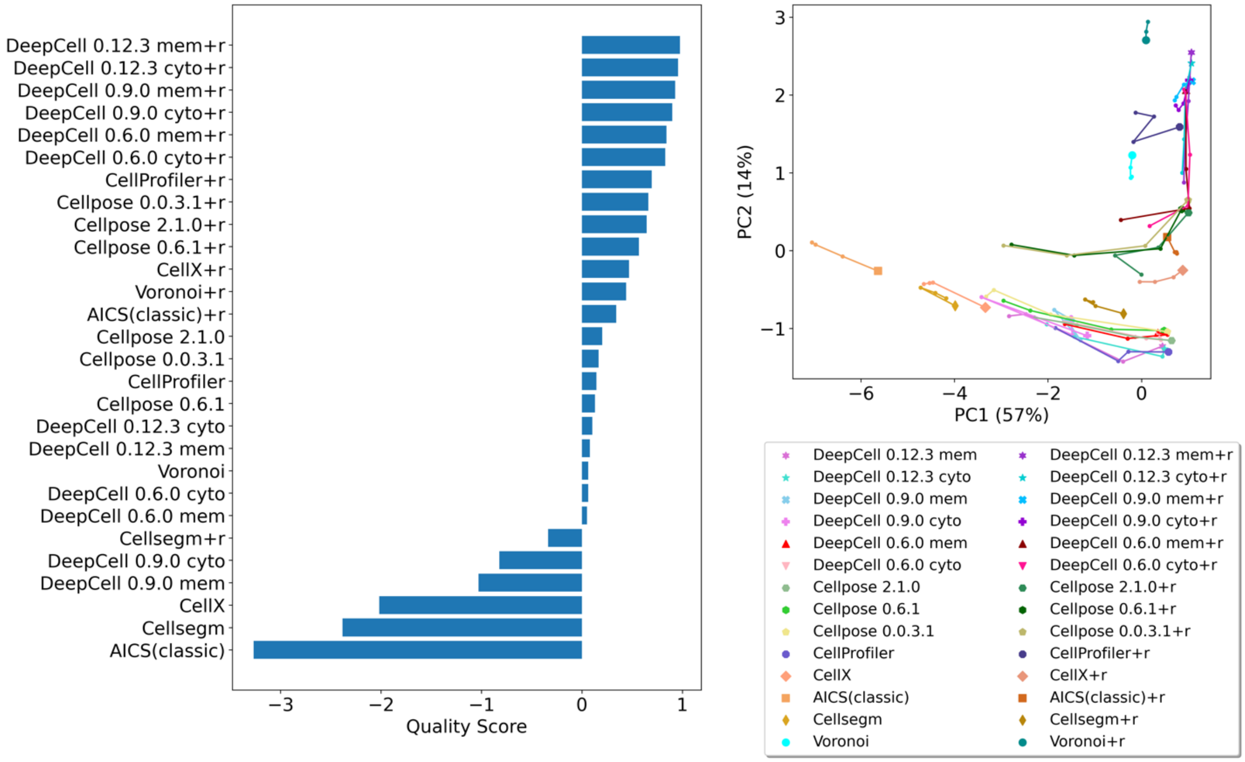


Supplementary Figure 12 Ranking of segmentation quality scores of all methods on Thymus images in CODEX and IMC datasets.

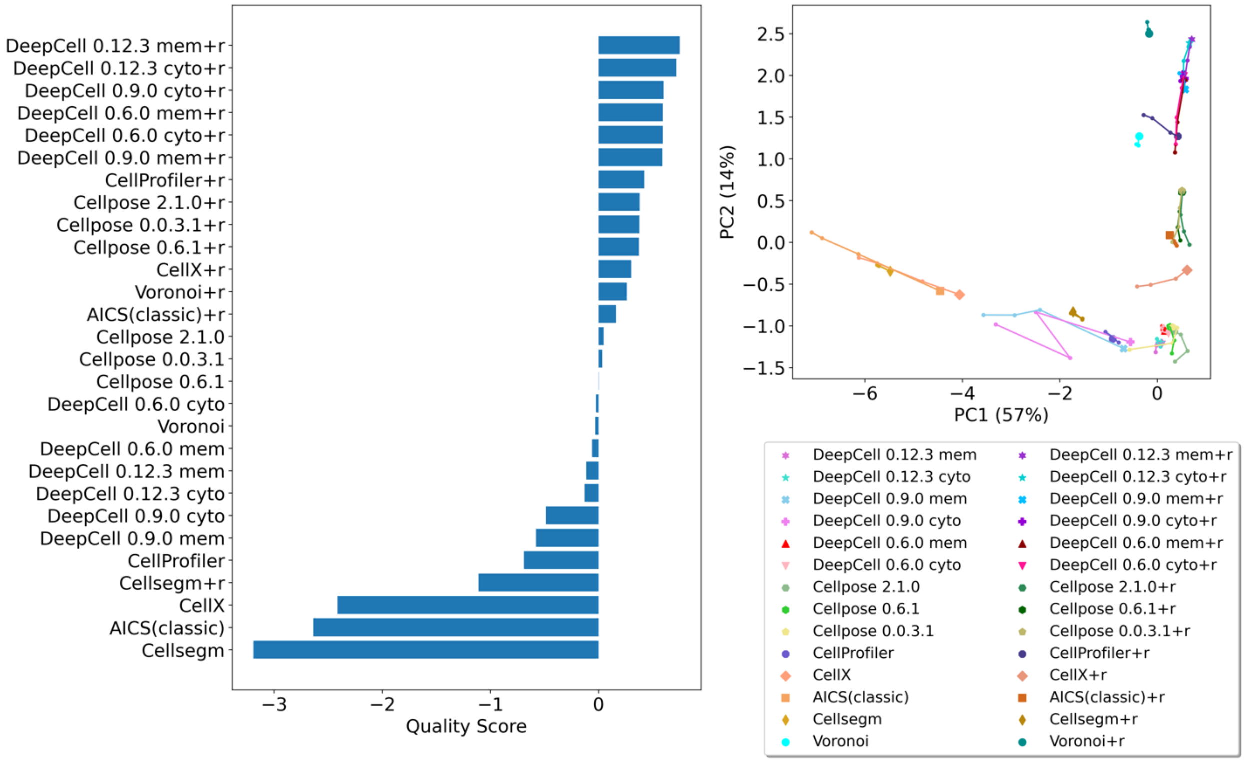


Supplementary Figure 13 Ranking of segmentation quality scores of all methods on Lymph Node images in CODEX and IMC datasets.

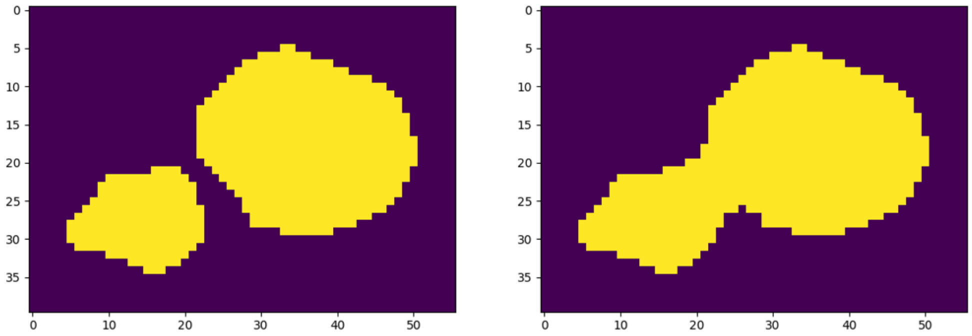


Supplementary Figure 14 Merging nuclei in simulating under-segmentation error. After merging contacting cells, we merged two separated nuclei (left panel) by operating morphological opening with a disk kernel (with tunable diameter) on the binary inverted mask of left panel to connect the gap between nuclei. The right panel shows final merged nuclei.

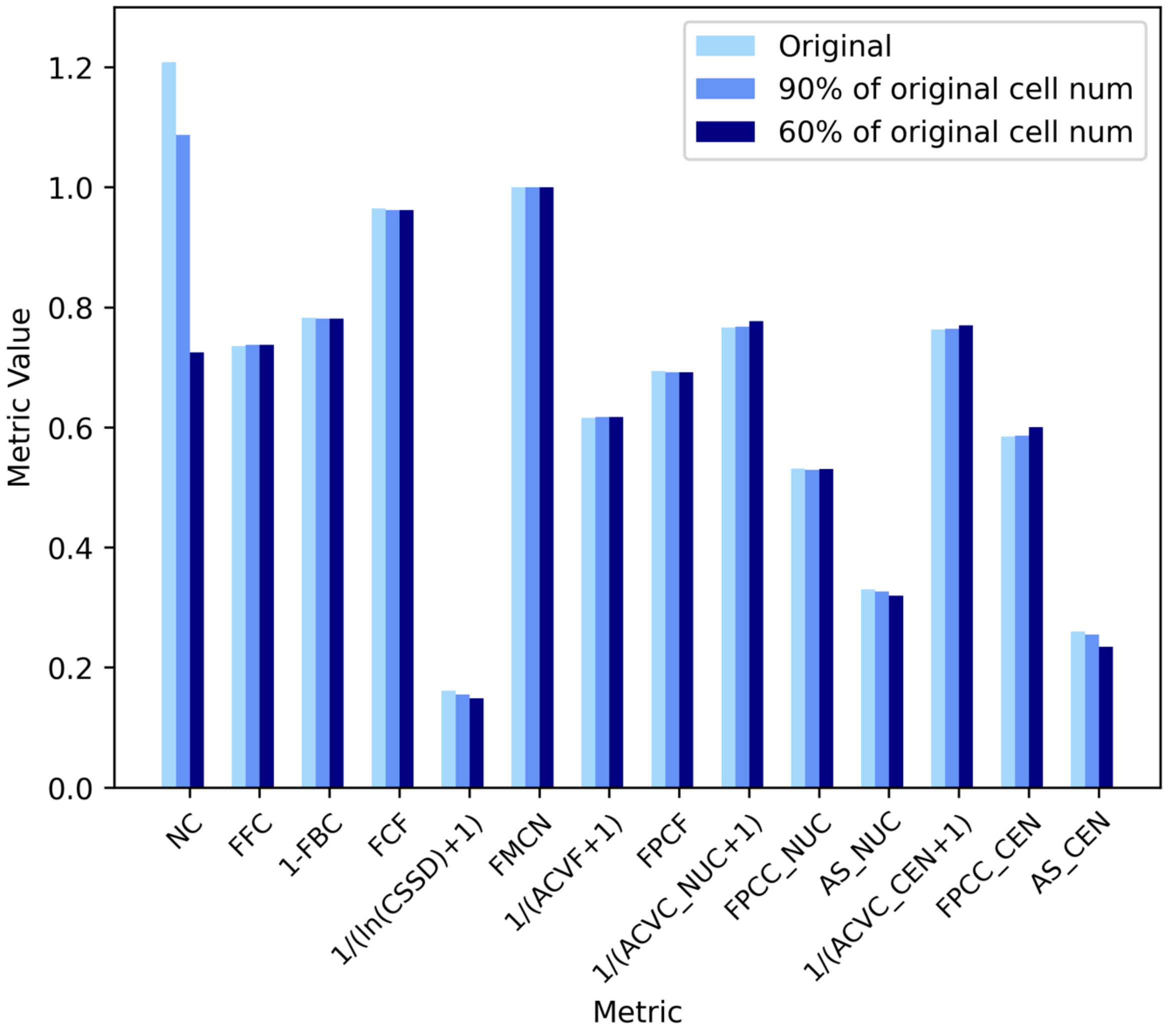


Supplementary Figure 15 The breakdown of metrics of simulated under-segmented masks. The density metric (NC) decreases with more severe under-segmentation. Cell shape uniformity metric is also monotonically decrease from original to the merged masks that have 60% of original cell number. Cell uniformity metrics (ACVC, FPCC, AS) have little change because new cell types that were introduced by merging contacting cells have high cell uniformity.

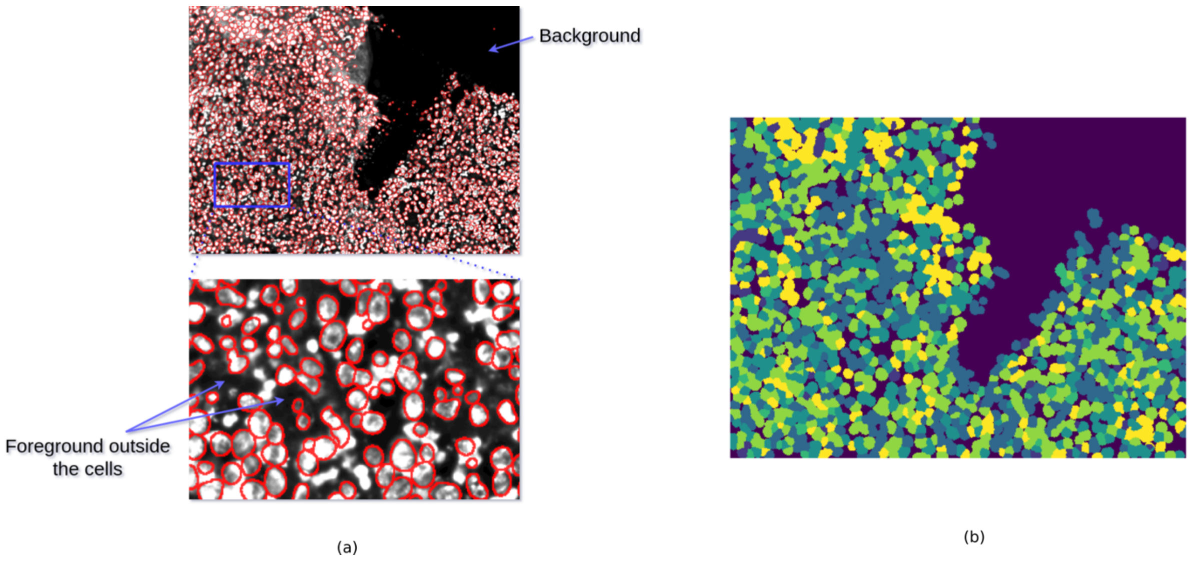


Supplementary Figure 16. Illustration of evaluation metrics. (a). This image comes from the sum of intensities of nucleus (DAPI) and cell membrane (E-cadherin) channels of tile R001_X001_Y001 of CODEX image HBM337.FSXL.564 which is of spleen tissue. The red contours are from the segmentation result of DeepCell 0.9.0 with cell membrane input on this image. (b). Clustered by the mean intensities per cell. We calculated the cell uniformity metrics within each cluster and averaged them across clusters.

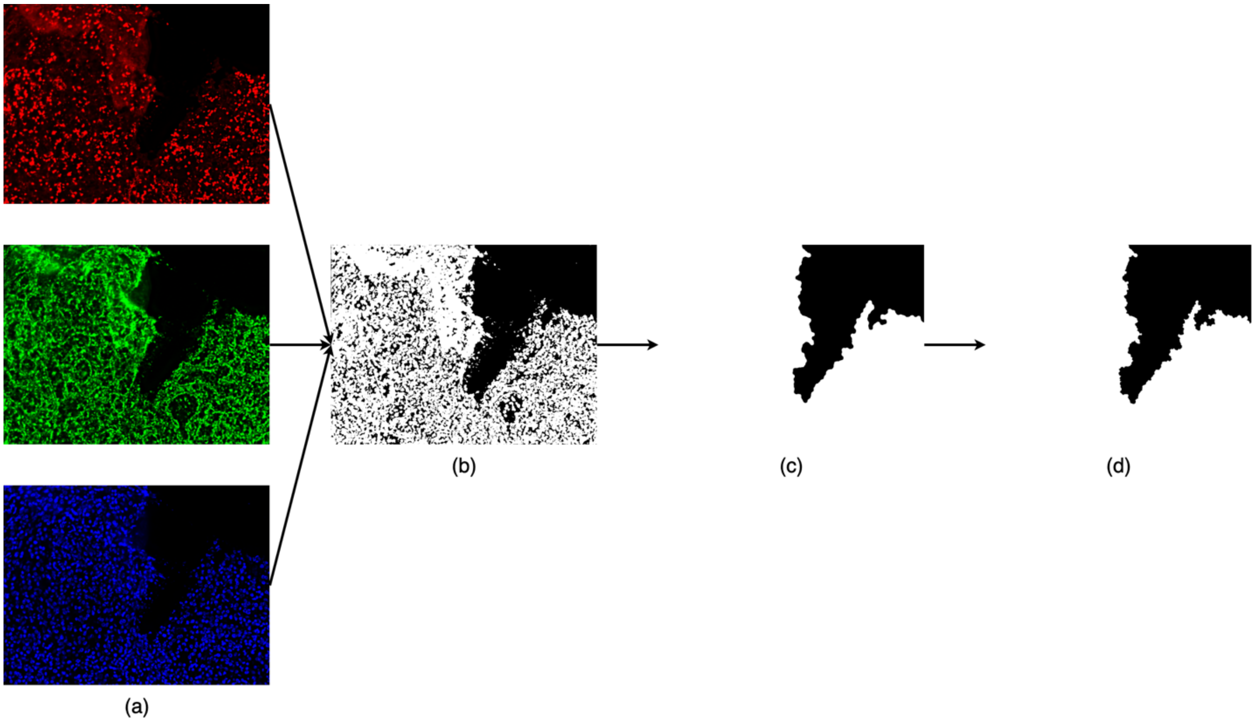


Supplementary Figure 17. Foreground-background segmentation. Mean thresholding was applied on the cell membrane (red, E-cadherin), cytoplasm (green, CD107a), and nuclear channels (blue, DAPI) images to draw the approximate outlines of tissue (a). The union of these binary images was obtained (b) and iterative morphological closing was applied to close the small and large gaps within the tissue (c). Lastly, a geodesic active contour was applied with background areas as seeds to correct the overly rounded boundaries between foreground and background (d).
